# Targeting neoantigens conserved across organs and species overcomes tumor immune escape

**DOI:** 10.1101/2025.09.25.678519

**Authors:** Guillaume Mestrallet, Ross W. Ward, Matthew Brown, Jesse Boumelha, Frederika Rentzeperis, Natalie Vaninov, Miriam Saffern, Ezekiel Olumuyide, Prerna Suri, Sreekumar Balan, Leandra Velazquez, Aparna Ananthanarayanan, Zhihong Chen, Aimee L. Lucas, Miriam Merad, Cansu Cimen Bozkus, Nicolas Vabret, Robert M. Samstein, Nina Bhardwaj

## Abstract

Neoantigen-targeted immunotherapies hold promise for cancer treatment, but current personalized approaches are time-consuming and costly. Here, we identify neoantigens encoded by *Ptprs* and *Igf2r* that are shared across murine mismatch repair-deficient colorectal and breast tumors and unexpectedly conserved in human colorectal, endometrial, gastric, and prostate cancers. These neoantigens elicit spontaneous, organ-spanning CD8+ T cell-mediated memory responses that are enhanced by immune checkpoint blockade. Vaccination with mRNA/lipid nanoparticles encoding these conserved neoantigens suppresses tumor growth across prophylactic and therapeutic models, including checkpoint-resistant orthotopic tumors. Tumor rejection is accompanied by antigen spreading, abscopal effects, and infiltration by clonally diverse T cells, dendritic cells, and MHC I/II+ macrophages producing CXCL9/10, CCL5/8, and TNF. Tumor cells also show activation of innate and adaptive pathways, including MHC and ISGs overexpression. Our results uncover a conserved anti-tumor immune mechanism and support the development of off-the-shelf neoantigen vaccines across tissues and species.

## Introduction

Immunotherapy has revolutionized cancer treatment by harnessing the power of the immune system to resist malignant cell growth. While mismatch repair deficient (MMRd) tumors often respond favorably to immunotherapy, a significant proportion of MMRd and most MMR proficient (MMRp) tumors exhibit resistance or relapse, underscoring the need for a deeper understanding of tumor immunogenicity and immune escape mechanisms (*1–7*). Tumor immunogenicity, the ability of tumors to elicit immune responses, is influenced by tumor-intrinsic factors, the tumor microenvironment (TME), and host immune responses. Due to an inability to repair DNA, MMRd tumors are characterized by an increased frequency of insertion/deletions (indels) that can encode neoantigens if they occur in coding regions (*1*). The overexpression of these neoantigens is accompanied by an inflamed TME that contributes to the clinical effectiveness of anti-PD-1 therapy in this patient population (*1*, *7*, *8*). However, tumors evade immune recognition through the upregulation of multiple immune checkpoints, clonal selection and myeloid immunosuppressive programs, enabling progression and therapeutic failure (*1*, *9–12*).

We previously discovered that T cells can spontaneously recognize highly immunogenic, frameshift-derived neoantigens shared across MMRd tumors (*13*), and recent studies suggest that neoantigen vaccination directed towards such targets can elicit protective responses in both murine models and patients (*14–24*). However, identifying neoantigens that are broadly shared among patients, which are simultaneously immunogenic and effective across histologies remains a major challenge for the development of broadly effective cancer immunotherapies (*25*, *26*). Achieving robust and durable anti-tumor responses in most patients remais a major challenge and will require systematic discovery of shared neoantigens and thorough evaluation of their immunogenicity across multiple cancer types and in relevant preclinical models (*15*). While clinical studies are assessing the efficacy of vaccination against shared neoantigens within *KRAS*, *p53* genes and MMRd frameshift mutations (*16*, *27*), it remains unclear whether this approach can elicit long-lasting immune memory responses that are conserved across tissues, which immune mechanisms sustain such responses and if they can overcome resistance to immune checkpoint blockade (ICB). It is also crucial to identify neoantigens shared across species to verify their immunogenicity in preclinical models before patient vaccination, and ultimately improve vaccination efficacy and safety.

In this study, we addressed these knowledge gaps by first screening for shared neoantigens across multiple MMRd tumors in murine models and in databases of human MMRd tumors across multiple histologies. We hypothesized that shared neoantigens contribute to both tumor-specific and cross-tumor immune memory, and thus represent promising targets for immunotherapy. Using integrated genomic analyses in murine tumor models, we identified new neoantigens shared across tumor types. We demonstrated that these shared frameshift-derived neoantigens are conserved not only amongst murine tumors but also within human tumors. Moreover, they can spontaneously elicit CD8+ T cell memory responses, which can be enhanced through combination of vaccination and ICB. We showed that vaccination against these conserved neoantigens overcomes resistance to ICB through neoantigen spreading and an abscopal effect, and induces the activation of broad innate and adaptive immune responses.

Together, our findings provide mechanistic insights into tumor immunogenicity and immune memory, and point to the translational potential of targeting neoantigens shared across organs and species for broad-spectrum cancer immunotherapy.

## Results

### Mutations and candidate neoantigens are shared between MMRd murine and human tumors in multiple organs

As frameshift (FS)-derived neoantigens are shared across MMRd patients (*13*), we hypothesized that murine MMRd tumors would similarly share frameshift mutations, and consequently neoantigenic epitopes which could be evaluated for off the shelf vaccine purposes. We first performed whole exome sequencing (WES) of colorectal CT26 and breast 4T1 tumors that were either wild type (WT) or had a deletion in the MMR gene MSH2, resulting in an MMRd phenotype. As expected, we observed more indels and single nucleotide variants (SNVs) in the MMRd vs WT tumors (**Figure 1 A, B**). Interestingly, CT26 tumors had high numbers of SNVs regardless of their MSH2 status, possibly accounting for their greater intrinsic immunogenicity (*8*). Notably, 4 indels (*Ptprs*, *Igf2r*, *Chrnb2* and *Vmn2r92*) and 45 SNVs were shared between CT26 and 4T1 MMRd tumors (**Figure 1 C**). All indel mutations shared by both MMRd cell lines are characterized by the loss of one nucleotide in a series of more than 7 identical nucleotides, with the majority (*Ptprs*, *Igf2r* and *Vmn2r92*) on chromosome 17, which happens to be the location of the *Msh2* gene. Similarly, when interrogating the cBioPortal CHORD database (*28*) of patients with a mutation in *MSH2*, we found that the most common shared frameshift mutations are on chromosome 17 (*p53*, *RNF43* and *PTPRS* genes), suggesting that MSH2 is key to repair these mutations. In addition, 50 indels and 2506 SNVs are shared by CT26 WT and MMRd tumors, 47 SNVs are shared by 4T1 WT and CT26 MMRd tumors (**Figure 1 C**). Finally, we also tested WT C11 lung tumor cells and found no sharing of indels or SNVs between CT26 MMRd and C11 tumors, a model that will be used as a control with no shared mutations in this study (**Figure 1 C**). Altogether, we identified new indels and SNVs that were shared between MMRd murine tumors in multiple cancer types.

**Figure 1.**
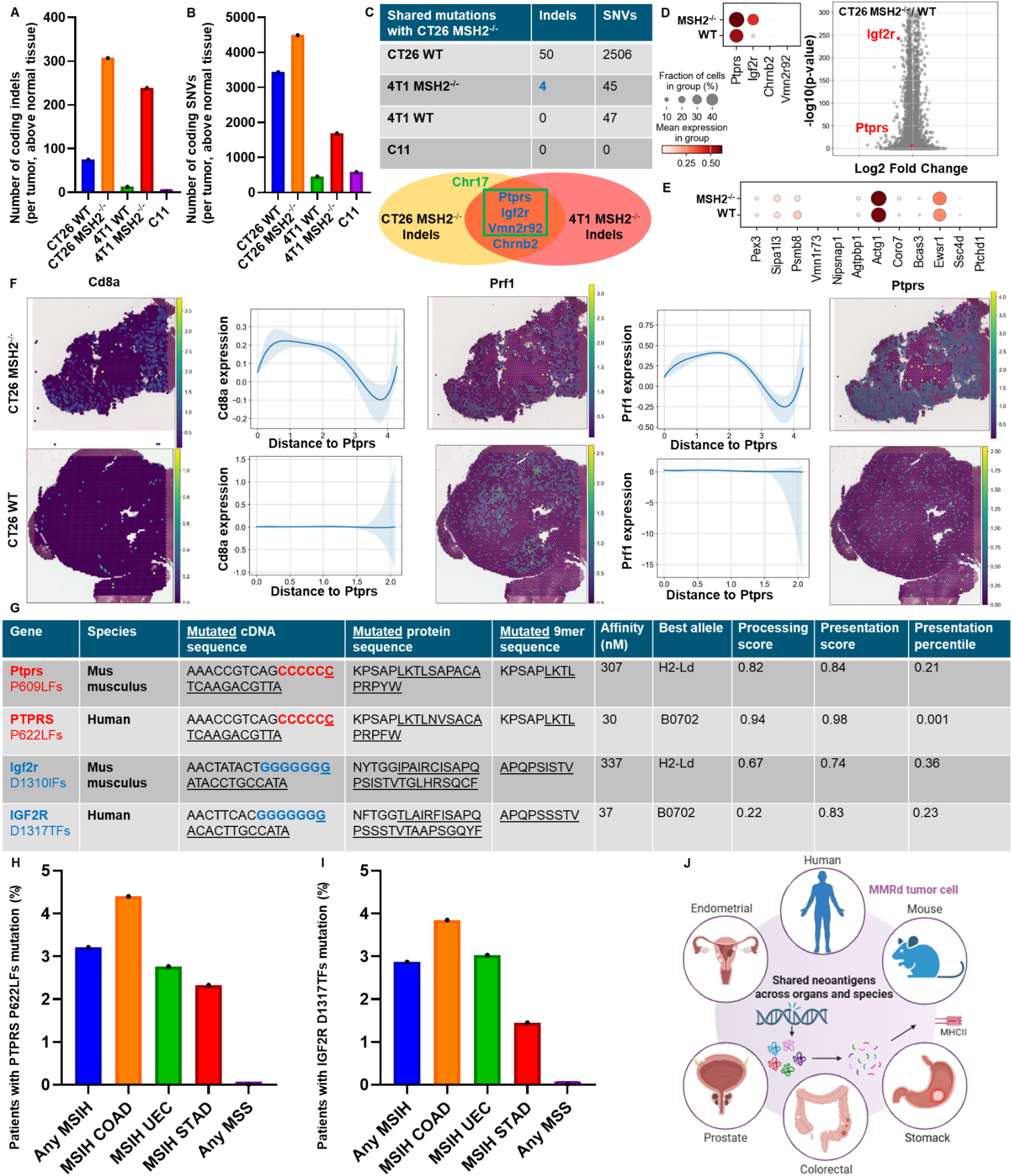
Mutations and candidate neoantigens are shared between MMRd murine and human tumors in multiple organs. Using whole exome sequencing, strelka, mutec2 and varcode, we measured the number of indel and snv mutations modifying the coding regions of genes in the CT26, C11 and 4T1 WT and MMRd cell lines ex vivo before tumor inoculation and after 28 days of tumor growth in vivo. 4 tumors were pooled together for each condition before WES. **A** Comparison of the number of indel mutations modifying the coding regions compared to normal tissue (spleen). **B** Comparison of the number of snv mutations modifying the coding regions compared to normal tissue (spleen). **C** Number of indel and snv mutations modifying the coding regions shared between each cell line and the CT26 MMRd cell line. **D** ScRNAseq of tumor cells was generated, following the exclusion of immune cells, in both CT26 WT and MMRd tumors. n=3 mice per group. ScRNAseq expression of genes encoding candidate MMRd neoantigens identified in table 1 by WES and MHCflurry. Wilcoxon test, p<0.05. **E** ScRNAseq expression of genes encoding candidate CT26 neoantigens identified in Table 2 by WES and MHCflurry. **F** SpatialRNAseq, expression of one of these genes encoding candidate MMRd neoantigens (*Ptprs*) in a CT26 MMRd tumor. Colocalization with Cd8+Prf1+ T cells was measured using scanpy and squidpy. N=3. **G** Sequences of the frameshift deletion mutations in *Ptprs* and *Igf2r* genes shared in both murine and human MSI-H tumors, with predicted affinity, best allele binding, processing score, presentation score and percentile by MHCflurry. **H** Percentage of MSI-H and MSS patients sharing *Ptprs* P622LFs mutation based on reanalyzed cbioportal data (*29*) with n = 623 MSI-H patients, including n = 318 colorectal cancer (COAD), n = 43 prostate cancer (PRAD) and n = 181 endometrial cancer (UEC). **I** Percentage of MSI-H and MSS patients sharing *Igf2r* D1317TFs mutation based on analysis of cbioportal TCGA data with n = 78 colorectal cancer (COAD), n = 69 stomach cancer (STAD) and n = 132 endometrial cancer (UEC). **J** Graphical summary of the identification of mutations and candidate neoantigens that are shared between MMRd murine and human tumors in multiple organs.

**Table 1.**
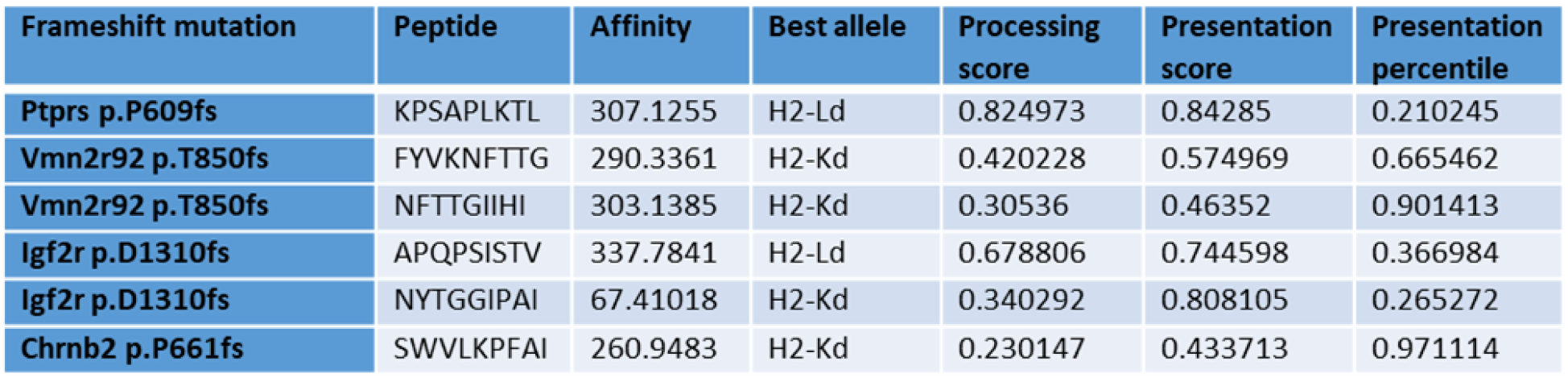
Peptides derived from indel mutations shared between 4T1 and CT26 MSH2 KO tumors with a predicted MHC I binding affinity below 500nM using MHCflurry. N=4 tumors pooled together for WES, N=4 cell lines growing pooled together for WES.

**Table 2.**
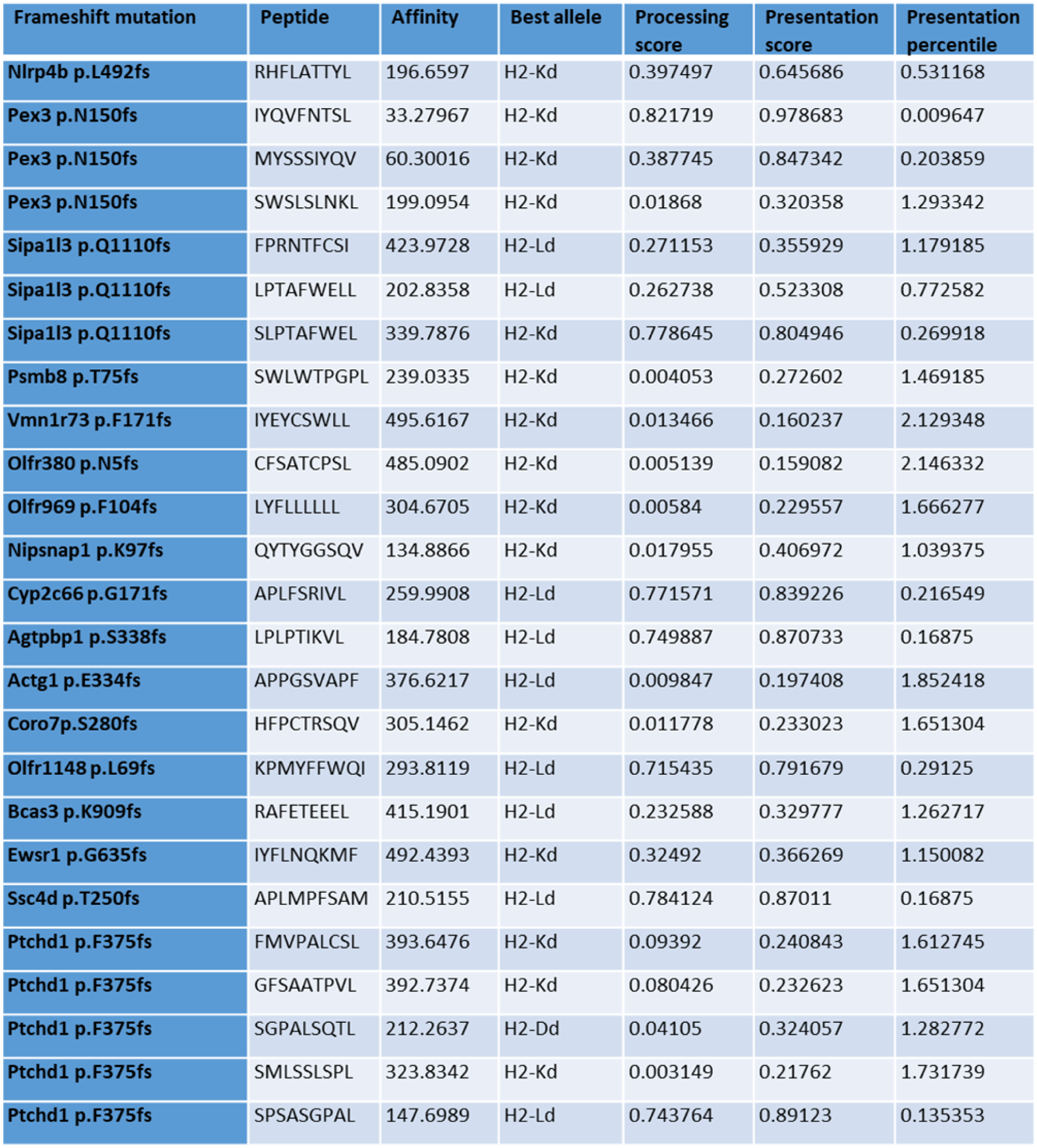
Peptides derived from indel mutations shared between CT26 tumors with a predicted MHC I binding affinity below 500nM using MHCflurry. N=4 tumors pooled together for WES, N=4 cell lines growing pooled together for WES.

**Table 3.**
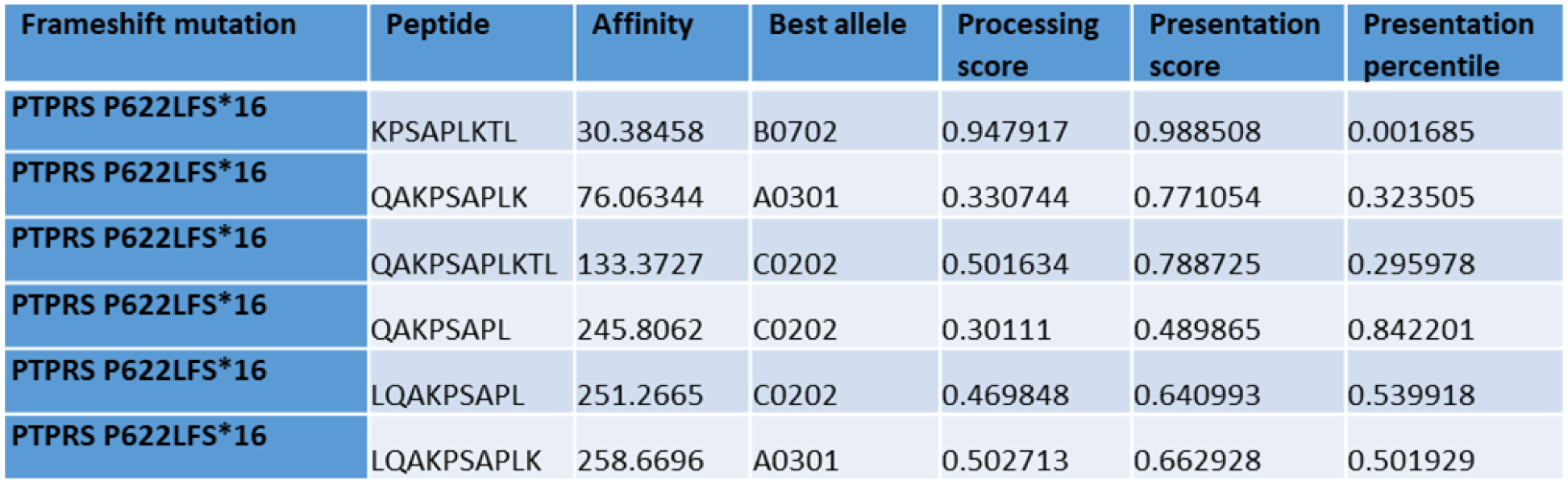
Peptides derived from Ptprs indel mutation shared between 4T1 and CT26 MSH2 KO tumors and human MSI-H CRC tumors with a predicted MHC I binding affinity below 500nM using MHCflurry. N=4 tumors pooled together for WES, N=4 cell lines growing pooled together for WES.

Given that CT26 and 4T1 MMRd tumors share indels, we speculated that they would give rise to shared sequences encoding potential immunogenic epitopes, as previously demonstrated for human CRC and endometrial cancers (*13*). We identified 6 peptide sequences (9 mers) derived from the 4 indel mutations in *Ptprs*, *Vmn2r92*, *Igf2r* and *Chrnb2* shared between 4T1 and CT26 MMRd tumors, that encoded epitopes with predicted binding affinity to H2-Kd and H2-Ld with IC50 values less than 500nM, using the MHCflurry prediction tool (**Table 1**). We also identified 25 MHC class I binding high affinity epitopes derived from indel mutations in the Nlrp4b, Pex3, Sipa1l3, Psmb8, Vmn1r73, Olfr380, Olfr969, Nipsnap1, Cyp2c66, Agtpbp1, Actg1, Coro7, Olfr1148, Bcas3, Ewsr1, Ssc4d and Ptchd1 genes which were solely present in CT26 WT and MMRd tumors but not present in 4T1 tumors (**Table 2**). Some peptides were also predicted to bind MHC II by applying netMHCII (**Table 4 and 5**). Our findings indicate that MMR deficiency in murine tumors increases mutational load and leads to shared indels and predicted neoantigenic epitopes, consistent with our observations in human tumors (*13*).

**Table 4.**
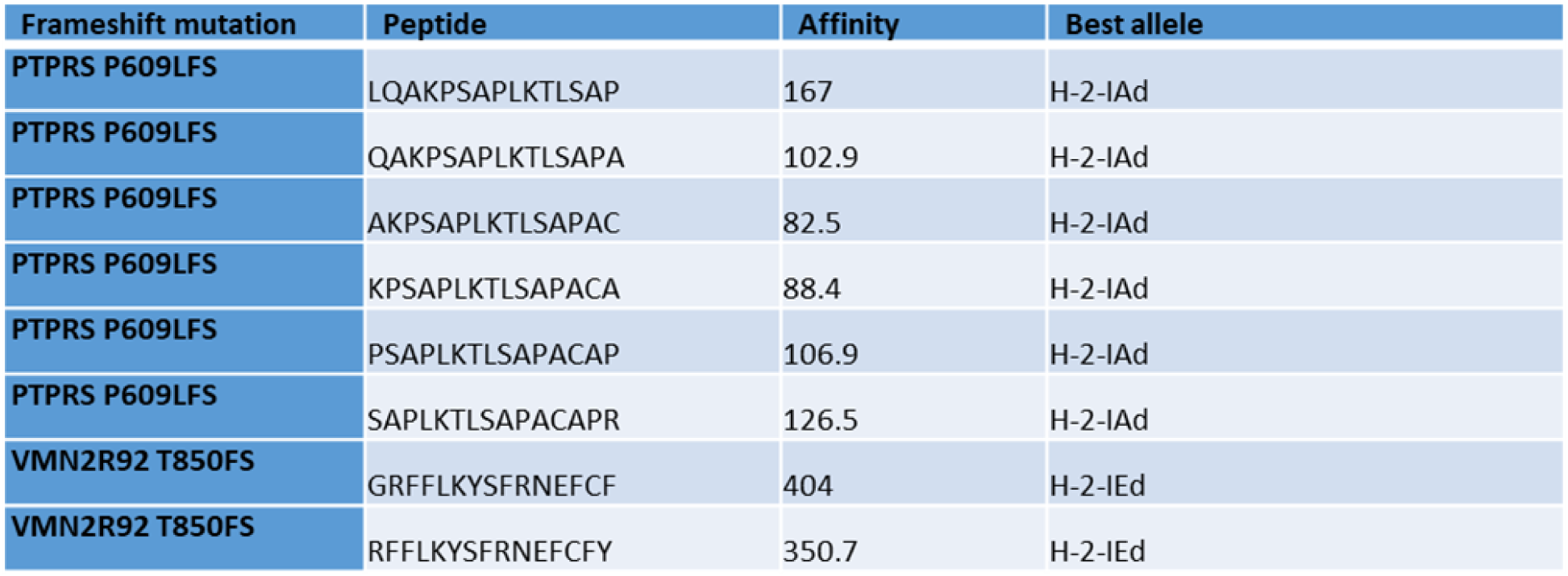
Peptides derived from indel mutations shared between 4T1 and CT26 MSH2 KO tumors with a predicted MHC II binding affinity below 500nM using netMHC II 2.3. N=4 tumors pooled together for WES, N=4 cell lines growing pooled together for WES.

The expression of the mutated *Ptprs* and *Igf2r* indel transcripts (**Table 1**) was verified by bulk RNAseq in CT26 MMRd tumors. We also undertook scRNAseq on 3 CT26 WT and 3 CT26 MMRd tumors and applied scanpy and leiden clustering, and confirmed the expression of *Ptprs* and *Igf2r* (**Figure 1 D**). We also observed the expression of several genes encoding potentially immunogenic candidate CT26 indel neoantigens (**Tables 2 and 5**) regardless of MSH status, including *Pex3, Sipa1l3, Psmb8, Agtpbp1, Actg1, Coro7, Bcas3, Ewsr1* and *Ssc4d* by scRNASeq (**Figure 1 E**). *Ptprs* and *Igf2r* WT transcripts are overexpressed in CT26 MMRd tumors (**Figure 1 D**), and we confirmed their expression in the whole CT26 MMRd tumors by spatial RNAseq (**Figure 1 F, Extended Figure 1**). We observed colocalization between *Cd8* and *Prf1* expression and genes encoding our candidate neoantigens in CT26 MMRd but not in WT tumors, indicating that their expression is associated with co-localization of cytotoxic T cells into the TME (**Figure 1 F, Extended Figure 1**). These findings confirmed the expression of transcripts encoding potential shared neoantigens in murine MMRd tumors, which may be eliciting specific T cell responses within the TME.

**Table 5.**
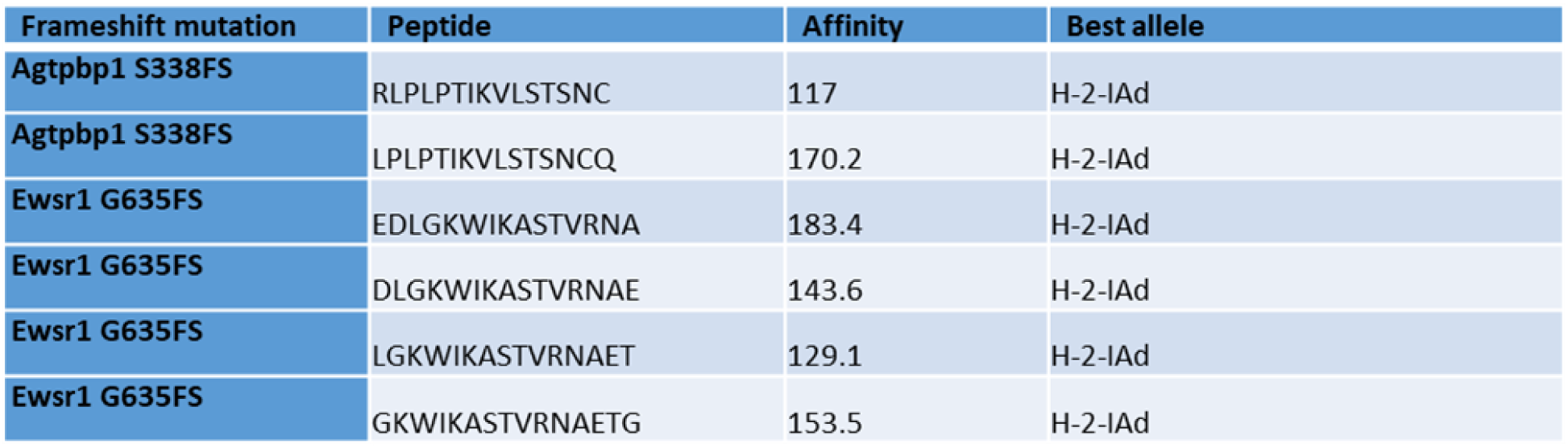
Peptides derived from indel mutations shared between CT26 tumors with a predicted MHC II binding affinity below 500nM using netMHC II 2.3. N=4 tumors pooled together for WES, N=4 cell lines growing pooled together for WES.

Finally, we investigated if these frameshift derived neoantigens shared by murine MMRd tumors might be relevant in human disease. We noted that the *Ptprs* P609 frameshift deletion mutation, identified in our murine tumors is also shared by 3.2% (e.g. 20 out of 623) of all patients with microsatellite unstable (MSI-H) tumors (**Figure 1 G, H**) and the predicted epitopes derived from this mutation are also predicted to bind human MHC I (**Table 3**) and MHC II (**Table 6**). This mutation is shared by 4.4% of MSI-H colorectal cancer patients, 2.8% of MSI-H endometrial cancer patients and 2.3% of MSI-H prostate cancer patients (**Figure 1 H**). Similarly, the *Igf2r* D1310 frameshift deletion mutation identified in our murine models is shared by 3.8% of MSI-H colorectal cancer patients, 3% of MSI-H endometrial cancer patients and 1.4% of MSI-H stomach cancer patients (**Figure 1 G, I**). Of note, this *Igf2r* indel was also found in one of our Lynch Syndrome CRC patients (**Extended Figure 2**). Moreover, we found that within individuals, FS indels are shared regardless of tumor origin, suggesting conserved immunogenicity across organs and emphasizing that incorporating these neoantigens as targets in vaccination approaches may yield broader impact (**Table 7**, **Extended Figure 2**). Overall, we identified new candidate FS indel-derived neoantigens shared across organs and conserved in murine and human MMRd tumors (**Figure 1 J**).

**Table 6.**
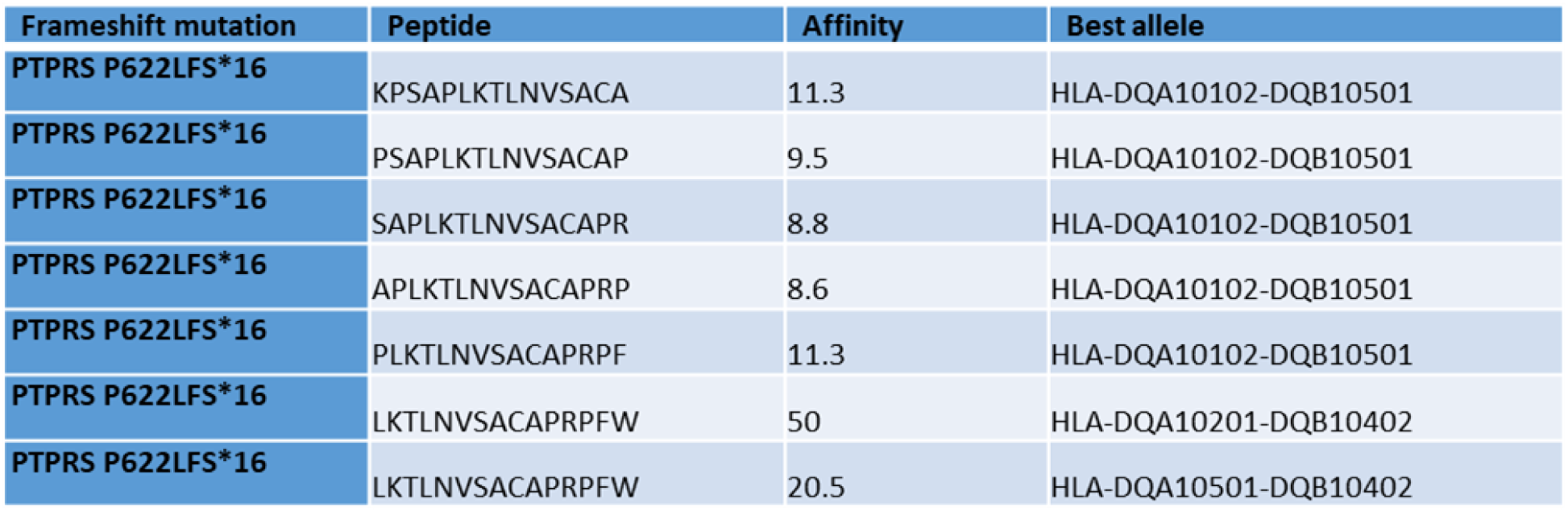
Peptides derived from Ptprs indel mutation shared between human MSI-H CRC tumors with a predicted MHC II binding affinity below 500nM using netMHC II 2.3.

**Table 7.** Mutations shared by several primary tumors on the same patient. Mutations shared by several primary tumors on the same patient based on reanalyzed cbioportal MSK-CHORD data, N = 10 patients.

| MSIH patient id | 1 |  | 2 |  | 3 |  |
| --- | --- | --- | --- | --- | --- | --- |
| Tumor type | CRC | BRCA | CRC | PRAD | PDAC | PRAD |
| JAK1 FS del K860Nfs*16 | **yes** | **yes** | no | no | no | no |
| MSH3 FS del K383Rfs*32 | no | no | **yes** | **yes** | no | no |
| KMT2D missense R3536H | no | no | no | no | **yes** | **yes** |

### Anti-tumor immune memory is conserved across organs and targets shared neoantigens through CD8+ T cells

Given our data showing the presence of neoantigens shared across organs and species, we investigated if these epitopes might induce a conserved and effective anti-tumor memory immunity. To do so, we first inoculated mice with CT26 MMRd tumors, and if the tumor was rejected, we assessed whether they would reject a second tumor of the same or different type. We observed that the primary tumor engraftment rate seems to negatively correlate with the number of mutations for each tumor type (**Figure 2 A**). These data are in accordance with our previous studies suggesting that a high tumor mutational burden (TMB) may be associated with increased anti-tumor immune responses (*7*, *8*).

**Figure 2.**
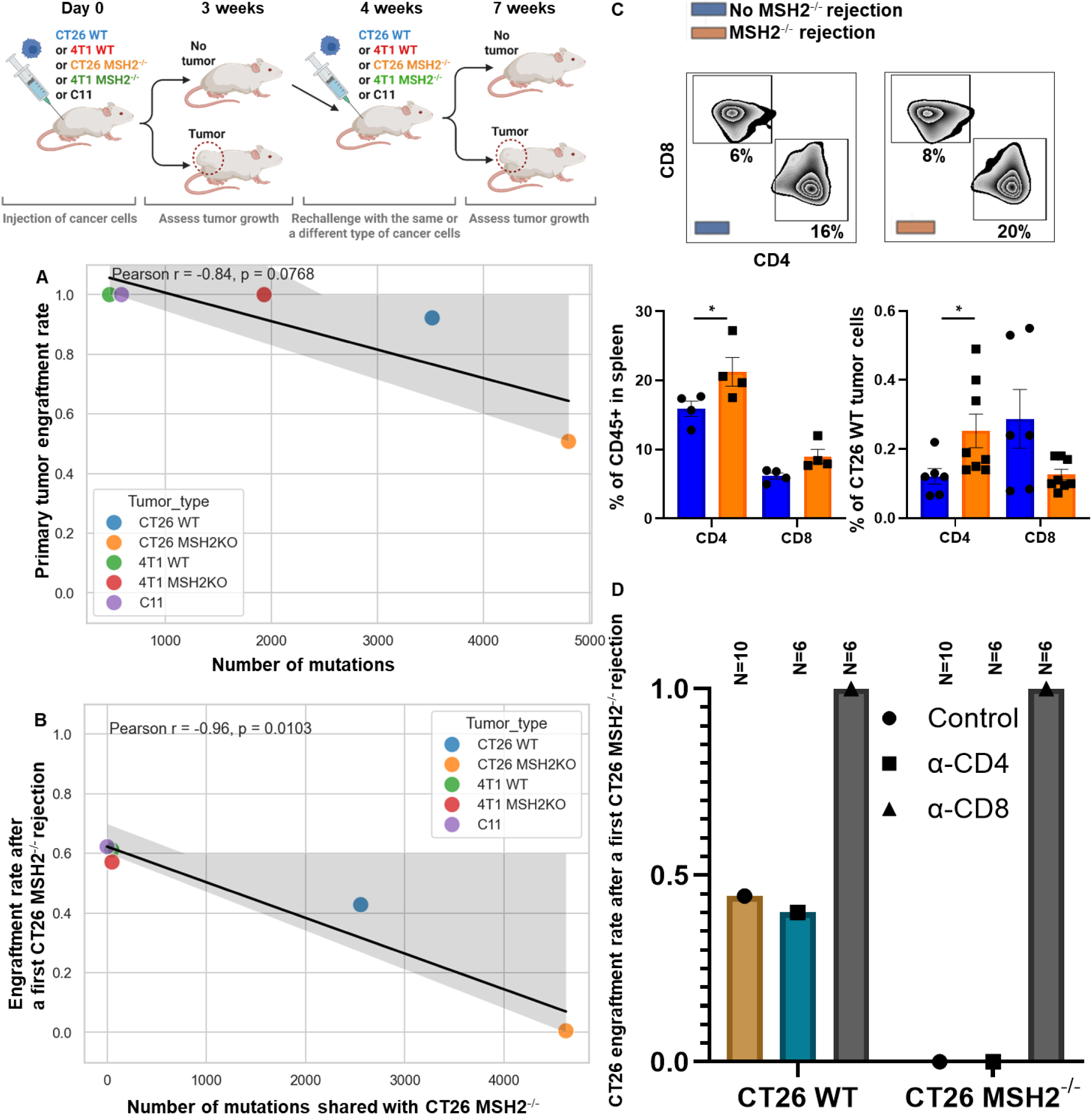
Anti-tumor immune memory is conserved across organs and targets shared neoantigens through CD8+ T cells. **A** Tumor engraftment rate of CT26, C11 and 4T1 WT and MMRd tumors (200,000 cells/tumor) and correlation with the number of mutations (Pearson correlation). N = 10 to 165. **B** Tumor engraftment rate of CT26, C11 and 4T1 WT and MMRd tumors (200,000 cells/tumor) following a first CT26 MMRd rejection and correlation with the number of mutations shared with the rejected CT26 MMRd tumor (Pearson correlation). N = 10. **C** Spleen composition from mice that rejected or not a first CT26 MMRd tumor (flow cytometry). CT26 WT tumor immune infiltration from mice that rejected or not a first CT26 MMRd tumor (flow cytometry). N = 4. Data are represented as mean ± SEM. **D** Anti-CD4 or anti-CD8 antibodies were used to deplete these populations before tumor rechallenge (100ug per mice for 4 days after a first CT26 MMRd tumor rejection and then once a week). N = 4 to 10. Data are represented as mean ± SEM.

As CT26 MMRd tumors had a 50% non-engraftment rate, the highest rate among all tumor types tested, we chose to implant a second tumor into these tumor-free mice. In all cases there was reduced engraftment of the second tumor regardless of its origin (**Figure 2 A, B**). Specifically, after a first rejection of CT26 MMRd tumors, we observed a 60% engraftment rate for 4T1 MMRd (49 shared mutations), for 4T1 (47 shared mutations) and for C11 (0 shared mutations), a 44 % engraftment rate for CT26 (2,556 shared mutations) and a 0% engraftment rate for MMRd CT26 tumors (4,624 shared mutations) (**Figure 2 B**). Importantly, the engraftment rate of a different tumor negatively correlated with the number of mutations shared with the first CT26 MMRd tumor (**Figure 2 B**, Pearson correlation). The reduction in tumor engraftment rates suggests that shared anti-tumor memory immune responses might be mediating the effect. Interestingly, even for the C11 that has no shared mutations with CT26 MMRd, the engraftment rate decreased compared to the baseline (100% vs 60%; **Figures 2 A, B**). To evaluate whether unresolved systemic inflammation induced by the primary tumor inoculation could impact secondary tumor growth, we waited 6 weeks, instead of 1, after the CT26 MMRd rejection before rechallenging the mice with C11 cells. We observed that engraftment rate increased from 60% to 100%, suggesting that inflammation and innate immunity were probably mediating the initial anti-tumor memory (**Extended Figure 2**). Overall, anti-tumor immune memory responses can be shared across cancer types, and they correlate with the presence of a higher number of neoantigens shared by the primary and the secondary tumors.

Based on the correlation between reduced secondary tumor engraftment rates following CT26 MMRd tumor rejection and shared mutation loads across tumors, we hypothesized that anti-tumor memory responses against conserved neoantigens may be mediated by T cells. We observed that spleens of mice that rejected CT26 MMRd tumors contained more CD4 + T cells (**Figure 2 C, Extended Figure 1**). Similarly, more CD4+ T cells and myeloid cells infiltrated secondary CT26 WT tumors following a first CT26 MMRd rejection and rechallenge compared to primary established CT26 WT tumors (**Figure 2 C, Extended Figure 3**). However, when depleting CD4+ or CD8+ T cells before rechallenge with a second MMRd tumor, we observed that CD4+ depletion did not change the percentage of engraftment, while CD8+ T cell depletion led to a 100% engraftment rate, indicating that specific and shared anti-tumor immune memory is primarily driven by CD8+ T cells (**Figure 2 D**). We observed the same CD8+ T cell dependency when rechallenge was with CT26 WT cells, meaning that both tumor-specific and shared anti-tumor immune memory are driven by CD8+ T cells. Altogether, we show that anti-tumor immune memory responses can be conserved across tumor types, and target shared neoantigens through CD8+ T cells.

### Targeting neoantigens conserved across human and murine tumors delays tumor growth and induces an abscopal effect

As anti-tumor shared memory is mainly mediated by CD8+ T cells (**Figure 2**), we hypothesized that the antigens targeted were primarily neoantigens shared across tumor types. We therefore tested whether vaccination against these neoantigens could modulate tumor growth. First, we designed an RNA-based vaccine encoding all the shared neoantigens that we identified in murine and human MMRd tumors (**Table 1**). mRNA lipid nanoparticles (LNPs) were produced as previously described (*30*, *31*), and characterization of antigen presentation after transfection of mRNA vaccine into the B16F10 cancer cell line was verified using OVA (**Extended Figure 2**). In prophylactic studies, we immunized each mouse with 5 μg of mRNA-LNPs diluted in 50 μL 1x PBS via retro-orbital injection at day 0. A booster dose was given on day 5 and PBS was used as a negative control. Mice were then challenged with tumors on day 10. Prophylactic vaccination with mRNA vaccine encoding shared neoantigens limited tumor growth in CT26 MMRd models (**Figure 3 A**), but immune escape occurred in the 4T1 MMRd model after 14 days. As expected, vaccination was not effective in CT26 and 4T1 WT models which do not express these MMRd associated and shared neoantigens. Thus, vaccination targeting neoantigens shared by human and murine MMRd tumors prevents tumor growth in a murine colorectal tumor model, though not in the breast cancer model.

**Figure 3.**
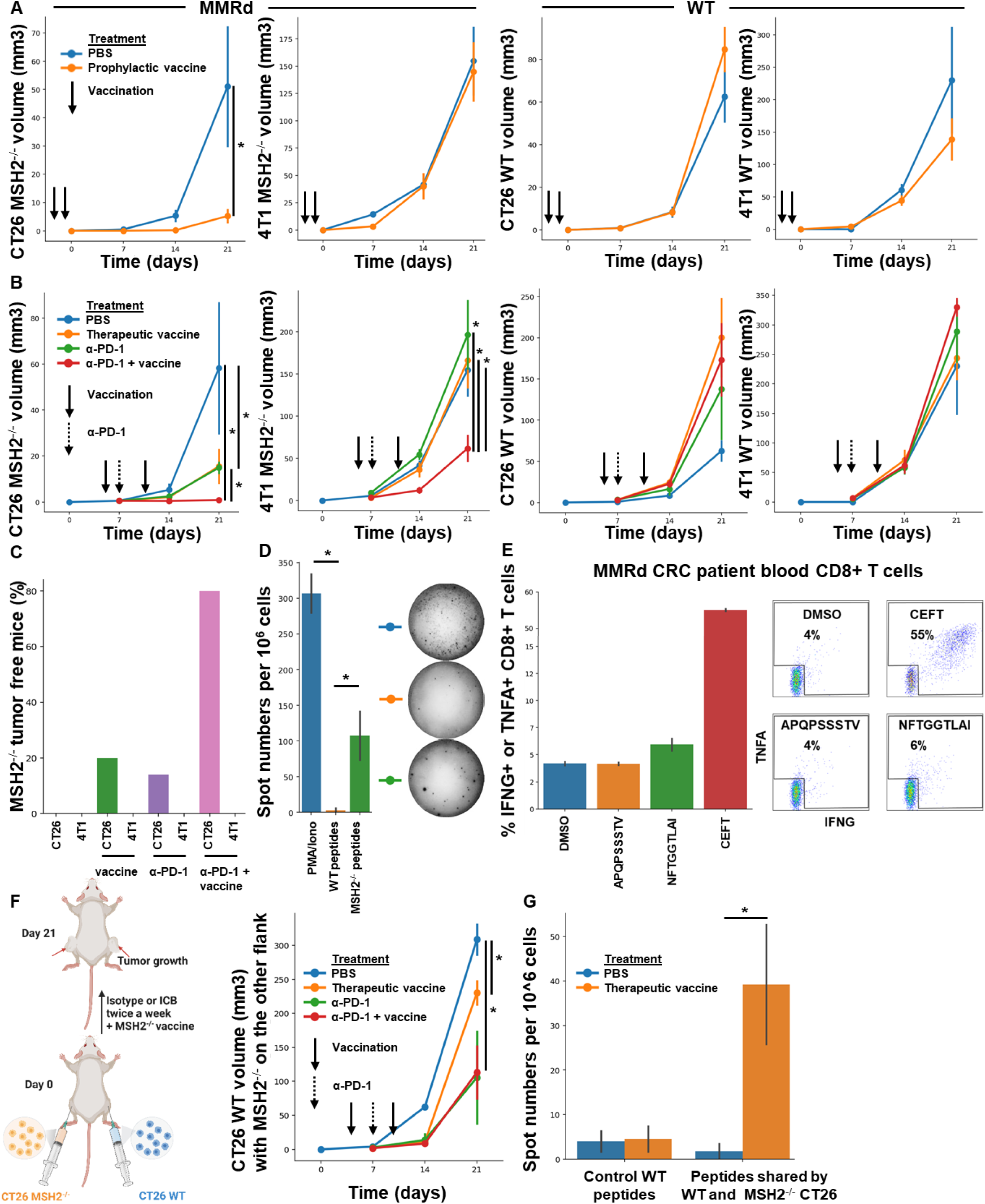
Targeting neoantigens shared by human and murine tumors across multiple organs delays tumor growth and induces an abscopal effect. **A/B** Tumor volume of CT26 and 4T1 WT and MMRd tumors (200,000 cells/tumor) in vivo. N = 5 to 10, Mann-Whitney U Test, pvalue<0.05. Data are represented as mean ± SEM. **A** Tumor volume of CT26 MMRd, 4T1 MMRd, CT26 WT and 4T1 WT tumors according to the presence of mRNA vaccine encoding shared MMRd neoantigens (5 μg mRNA-LNPs 10 days and 5 days before tumor inoculation). **B** Tumor volume of CT26 MMRd, 4T1 MMRd, CT26 WT and 4T1 WT tumors according to the presence of anti-PD-1 (100 ug twice a week 7 days after tumor inoculation) and mRNA vaccine encoding shared MMRd neoantigens (5 μg mRNA-LNPs at 5 days and 10 days after tumor inoculation). PBS was used as a negative control. **C** Percentage of CT26 and 4T1 MMRd tumor complete elimination after ICB and therapeutic vaccination combinations (%). Vaccination efficacy in orthotopic settings for both models was also confirmed in **Extended Figure 2**. **D** Elispot assay was performed with blood cells from vaccinated mice and stimulation with WT or mutated peptides shared by MMRd tumors and predicted to be immunogenic identified in Table 1. N=4, Mann-Whitney U Test, pvalue<0.05. **E** Intracellular cytokine staining of CD8+ T cells following stimulation of blood T cells from one MMRd CRC patient with the Igf2r indel mutation with fs-neoantigen peptides (patient data and sequences are in **Extended Figure 2**). DMSO as negative control, CEFT as positive control. N=2. Data are represented as mean ± SEM. **F** CT26 WT and CT26 MMRd tumors were inoculated on opposite flanks and mice were treated with anti-PD-1 (100 ug twice a week 7 days after tumor inoculation) and mRNA vaccine encoding shared MMRd neoantigens (5 μg mRNA-LNPs at 5 days and 10 days after tumor inoculation). Tumor volume of CT26 WT tumors. N= 5 to 10, Mann-Whitney U Test, p value<0.05. Data are represented as mean ± SEM. **G** Elispot assay was performed with blood cells from vaccinated mice with bilateral CT26 WT/MMRd tumors and stimulation with control WT or mutated peptides shared by CT26 tumors regardless MSH2 status and predicted to be immunogenic identified in Table 2. N=4, Mann-Whitney U Test, pvalue<0.05.

Next, we investigated if vaccination was effective in the therapeutic setting and could synergize with anti-PD-1 therapy. Mice were challenged with tumors on day 0, and immunized with mRNA-LNPs on day 5. A booster dose was given on day 10. 100µg of anti-PD-1 was administered intraperitoneally in 100 µl of PBS twice a week, after day 7. Therapeutic vaccination with mRNA vaccine encoding shared neoantigens + anti-PD1 limited tumor growth in both the CT26 and 4T1 MMRd orthotopic and subcutaneous models (**Figure 3 B, Extended Figure 2**), compared to untreated and anti-PD-1 only treated mice. VAF levels of *Ptprs* and *Igf2r genes* were only decreased following vaccination in the CT26 model, indicating a limited immunoediting in the 4T1 model (**Extended Figure 2**). As expected, vaccination was not effective in CT26 and 4T1 WT models that do not express these MMRd associated and shared neoantigens (**Figure 3 B**), and vaccination against irrelevant neoantigens did not delay MMRd tumor growth (**Extended Figure 2**). Moreover, 80% of the successfully engrafted CT26 MMRd tumors were eliminated following vaccination and anti-PD1 treatment, compared to 0% without treatment, 20% with vaccine alone and 14% with anti-PD1 alone (**Figure 3 C**). Overall, tumor growth is delayed by therapeutic mRNA vaccination targeting neoantigens shared by MMRd human and murine tumors, even in models resisting PD-1 blockade.

To confirm vaccine specificity, we performed elispot using blood PBMC from vaccinated mice and restimulated with the 6 MMRd neoantigens identified in **Table 1** or corresponding WT peptides. It confirmed that the vaccines elicited immunity towards these MMRd neoantigens shared by human and murine tumors and incorporated in our vaccine constructs (**Figure 3 D**). We also confirmed these results with a spheroid killing assay using the same settings (**Extended Figure 2 G**). As the *Igf2r* indel was also identified in one of our Lynch Syndrome MMRd CRC patients (**Extended Figure 2**), we determined whether the patient developed ex vivo spontaneous endogenous immunity towards neoepitopes encoded by this indel. Indeed, PBMC showed evidence of T cell immunity towards the class-I restricted NFTGGTLAI epitope compared to the DMSO control (**Figure 3 E**).

Finally, we hypothesized that vaccination targeting shared neoantigens may also induce epitope spreading where the immune response initially targeting the vaccine epitopes broadens and results in recognition of additional epitopes, leading to an abscopal effect. Vaccination with mRNA vaccine encoding shared MMRd neoantigens + anti-PD1 in the presence of a WT tumor on one flank and a MMRd tumor on the other flank resulted in reduced growth of the WT tumor in the presence of either anti-PD-1 +/- vaccination (**Figure 3 F)**. We hypothesized that this may be due to the recognition by T cells of other shared neoantigens between CT26 WT and MMRd tumors following treatment (**Figure 1 C, E, Table 2**). Thus, we performed elispot and spheroid killing assays with PBMCs from these vaccinated or non-vaccinated mice using neoantigen peptides listed in **Table 2** that are shared by CT26 tumors regardless of their MSH2 status and not incorporated in our vaccine. Vaccination and anti-PD-1 induced neoantigen spreading and increased abscopal tumor control (**Figure 3 G, Extended Figure 2 H**). To directly assess whether vaccination against MMRd neoantigens induces antigen spreading, we engineered CT26 MMRd cells to express GFP and monitored GFP-specific T cell responses using GFP-tetramer staining. Following vaccination, we detected GFP-tetramer-positive T cells, demonstrating that immunity extended to antigens not included in the vaccine (**Extended Figure 2**). We next investigated whether spreading could also be observed in vivo across tumors of different origins. Mice were bilaterally inoculated with CT26 and 4T1 WT or MMRd tumors. Even in the absence of vaccination, we observed evidence of abscopal effects and potential neoantigen spreading between tumors (**Extended Figure 3**). Importantly, these effects were amplified by combining multiple checkpoint inhibitors (anti-PD-1, anti-CTLA-4, anti-LAG3, and anti-TREM2) (*8*), suggesting that checkpoint blockade can broaden and potentiate antigen spreading across tumors of different origins (**Extended Figure 3**). Altogether, tumor growth is delayed by prophylactic and therapeutic mRNA vaccination targeting neoantigens shared by MMRd human and murine tumors, even in models resisting PD-1 blockade. It in turn allows the recognition of other shared neoantigens, and the abscopal tumor control of distant tumors that do not express the neoantigens initially targeted by vaccination.

### Targeting conserved neoantigens through vaccination promotes the recruitment of stem-like and proliferating CD4+ and CD8+ T cells, MHC+ macrophages, and innate responses

We investigated which immune cells mediate the response to vaccination against shared neoantigens. Following vaccination, 4T1 and CT26 tumors were more infiltrated by T cells, especially CD4+ and CD8+ subsets expressing higher levels of TCF, a marker of stem-like T cells (*32*), PD1 and CD69 (**Extended Figure 2)**. Of note, CT26 MMRd tumors were more infiltrated by CD8+ T cells compared to the less vaccine responsive 4T1 tumors. Thus, stem-like and proliferating CD4+ and CD8+ T cells may mediate response to vaccination.

scRNAseq of CT26 MMRd tumors revealed that vaccination led to the recruitment of MHC+ macrophages, DCs and CD4+ T cells, while PD-1 blockade was characterized by CD8+ T cell, C1Q+ macrophage infiltration, and a diminution of NK cells compared to untreated tumors (**Figure 4 A, B, Extended Figure 4**). The combination of vaccination and PD-1 blockade induced higher numbers of MHC I/II+ CXCL9+ CXCL10+ CCL5+ CCL8+ TNF+ macrophages, DCs, mast cells, CCL5+ CCL8+ NK cells, CD4+ and CD8+ T cells expressing TCF, Ki67 and CCL8 (**Figure 4 B, C, Extended Figure 4**). CCL5 and CCL8 are chemokines recruiting various immune cells such as T cells, eosinophils, basophils, monocytes, NK cells, and DCs (*33*, *34*), while CXCL9 and CXCL10 are chemokines that attracts CD8+ cytotoxic T cells (*35*). Vaccination also reduced infiltration by SPP1+ macrophages, previously described as immunosuppressive (*36*), monocytes and exhausted T cells (**Figure 4 B, C, Extended Figure 4**). Thus, mRNA/LNP vaccination promoted the recruitment of antigen presenting cells as well as CD4+ T cells, and acted in synergy with PD-1 blockade to promote an effective anti-tumor immune response.

**Figure 4.**
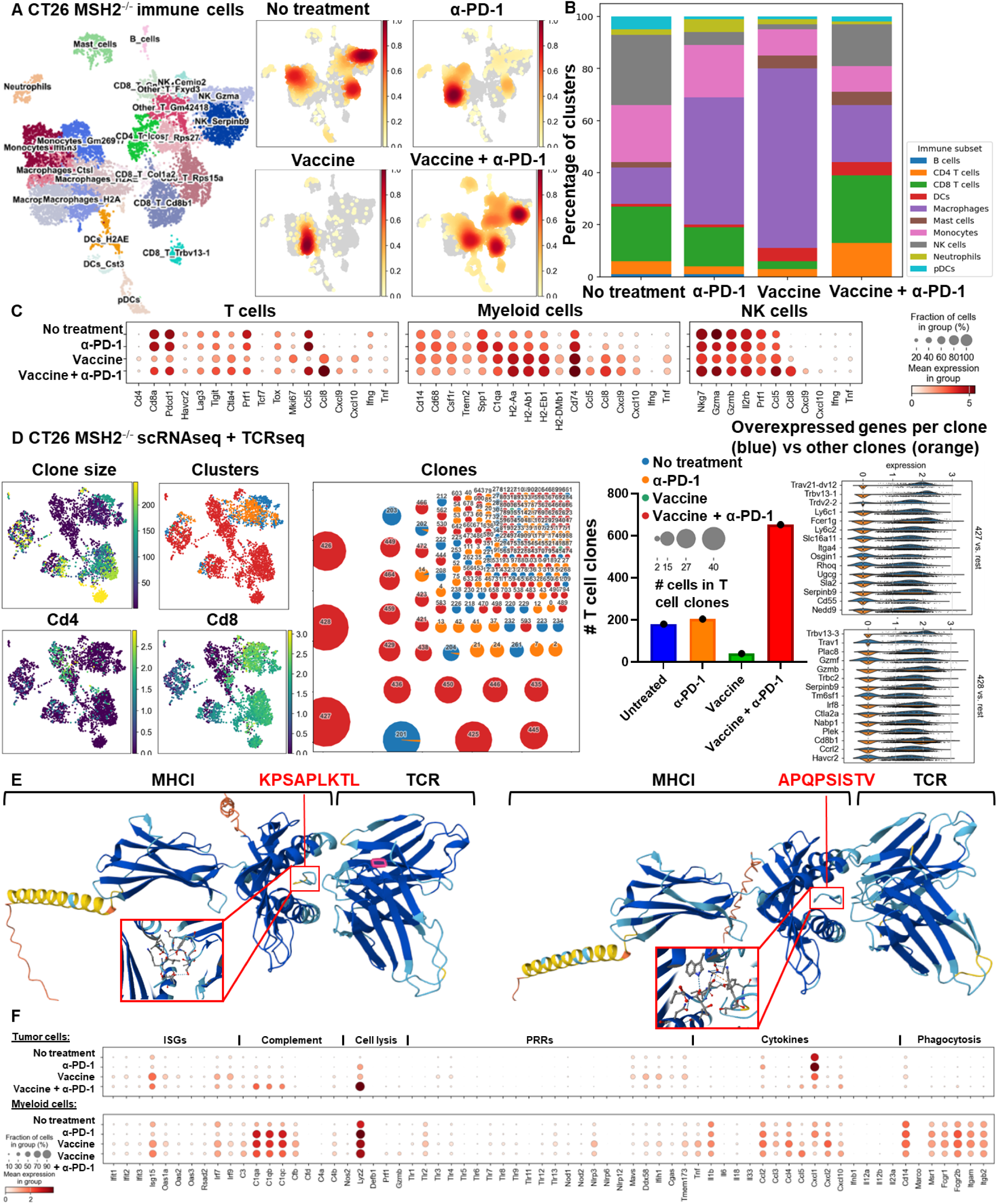
Targeting conserved neoantigens promotes the recruitment of MHC+ macrophages, stem-like and proliferating CD4+ and CD8+ T cells, and innate responses. **A** ScRNAseq UMAP of immune cells in CT26 MMRd tumors with or without anti-PD-1 therapy or vaccination encoding shared MMRd neoantigens (5 μg mRNA-LNPs at 5 days and 10 days after tumor inoculation). UMAP of immune clusters subsequent to Leiden clustering by scanpy. **B** Percentage of cells in each immune cluster. **C** Gene expression within each immune cluster. **D** ScTCRseq UMAP of immune cells in CT26 MMRd tumors with or without anti-PD-1 therapy or vaccination encoding shared MMRd neoantigens. UMAP of immune clusters subsequent to Leiden clustering by scanpy and clonal expansion of T cell clones. Identification of T cell clones, number of cells in each clonotype for each tumor type and gene expression within each main T cell clone (clone of interest : blue vs other clones : orange). **E** Alphafold prediction of the binding of the neoantigens shared across organs and species (KPSAPLKTL for *Ptprs*, APQPSISTV for *Igf2r*) with murine MHC I (H2Dd) and the TCRs from the most expended T cell clones in the tumors. **F** ScRNAseq of tumor and myeloid cells in CT26 MMRd tumors with or without anti-PD-1 therapy or vaccination encoding shared MMRd neoantigens (5 μg mRNA-LNPs at 5 days and 10 days after tumor inoculation). Expression of innate immune genes in tumor and myeloid cells.

We next determined whether vaccination promoted the clonal expansions of T cells. By applying TCRseq together with scRNAseq on CT26 MMRd tumor infiltrating cells, we observed expansion of multiple clones of CD4+ and CD8+ T cells following treatment with the combination of PD-1 blockade and vaccination compared to untreated or anti-PD-1 treated tumors (**Figure 4 D, Extended Figure 5**). A limited number of T cells was obtained for tumors treated with vaccination alone, limiting the analysis for this condition. The main expanded clones following vaccination and ICB combination expressed high levels of serpin genes encoding proteins known to inhibit serine proteases and that are key regulators of immune homeostasis (*37*). Using Alphafold, we predicted the binding between the neoantigens shared across organs and species (KPSAPLKTL for *Ptprs*, APQPSISTV for *Igf2r*) with murine MHC I (H2Dd) and the TCRs from the most expended T cell clones in the tumors (**Figure 4 E**). These neoantigens are predicted to bind MHC I and TCRs from the mostly expanded clones. Overall, vaccination targeting neoantigens shared across organs and species acts in synergy with PD-1 blockade to better control tumor growth. Through deep analysis, we show the recruitment of multiple CD4+ and CD8+ T cell clones, stem-like and proliferating T cells, NK cells, DCs and MHC+ antigen presenting macrophages expressing multiple cytokines.

Finally, we investigated if the expression of the genes encoding adaptive and innate immune responses by tumor cells was modified after therapy. By performing scRNAseq on CT26 MMRd tumor cells alone, we observed that the expression of *Ptprs* and *Igf2r* was significantly decreased after the combination of vaccination and PD-1 blockade compared to anti-PD-1 therapy alone (**Extended Figure 6**). On the other hand, the expression of genes involved in antigen processing and presentation via MHC, cytokine signaling, cell killing, interferon and humoral responses were increased in the tumor cells after vaccination compared to the baseline (**Extended Figure 6**). Specifically, vaccination promoted the activation of innate immune responses in tumor and myeloid cells, especially interferon-stimulated genes (ISGs) (*Isg15* ubiquitin-like modifier, *Interferon regulatory factor 7*, *Ifit1*, *Ifit2*, *Ifit3* – *Interferon-induced proteins*, *Oas1a*, *Oas2*, *Oas3* – *2’-5’-oligoadenylate synthetases* and *Rsad2* (*Viperin*)) (**Figure 4 F**). Activation of the complement system (*C3*, *C1q*, *Cfb*, *Cfd*, *C4b*, *C4a*) and cytolytic effectors (*Nitric oxide synthase*, *Lysozyme 2*, *Perforin* and *Gzmb*) were also observed after vaccination in tumor and myeloid cells, together with up regulation of pattern recognition receptor (PRR) (*RIG-1*, *CGAS-STING*, *MAVS*) and cytokine profiles (**Figure 4 F**). On the contrary, the expression of genes involved in glycolysis, tumor growth, metabolism was reduced in tumor cells compared to the untreated condition (**Extended Figure 6**). Thus, vaccination reduced the expression of genes involved in metabolic activity in tumor cells, while increasing the expression of genes related to adaptive and innate immune responses.

Altogether, vaccination against conserved neoantigens promoted infiltration of stem-like and proliferating CD4+ and CD8+ T cells, MHC+ macrophages and DCs, while reducing immunosuppressive macrophages and exhausted T cells. In combination with PD-1 blockade, vaccination enhanced clonal expansion of T cells, induced antigen presentation and interferon-stimulated gene expression in tumors, and synergistically improved anti-tumor immune responses.

## Discussion

Our study uncovers new insights into the mechanisms by which anti-tumor immune memory responses are both conserved and shared across tumor types by targeting neoantigens shared amongst MMRd tumors in murine models. Through an integrative analysis of genomic, transcriptomic, and immunologic data, we identified a set of frameshift-derived neoantigens that are not only recurrent across murine colorectal and breast MMRd tumors, but also conserved in human MMRd colorectal, endometrial, prostate, and gastric cancers. Tumor mutations in these genes were also found in other species such as dogs (*38–40*). These neoantigen sequences, such as those arising from recurrent FS mutations in *Ptprs* and *Igf2r*, were predicted to encode epitopes that bind both MHC I and MHC II molecules and were shown to be expressed in untreated tumor cells, co-localizing with CD8+ T cell infiltration and cytotoxic markers, suggesting their in vivo immunogenicity.

A key finding is that neoantigen-specific immune memory responses can be both tumor-specific and cross-protective, and that this cross-protection correlates with the number of shared mutations between tumors. Importantly, CD8+ T cells were essential for mediating this shared memory, underscoring their critical role in targeting conserved neoantigens across distinct tumors. These responses were sufficient to elicit abscopal effects, limiting the growth of distant tumors of different histologies within the same host, even in the context of immune checkpoint resistance.

We demonstrate that mRNA/LNP-based vaccination targeting these conserved neoantigens can prevent and control tumor growth in multiple MMRd tumor models, and that it acts synergistically with PD-1 blockade to overcome immune resistance. Notably, combination therapy not only enhanced local tumor control but also induced epitope spreading and abscopal tumor rejection, including of tumors not directly targeted by the vaccine. This supports the concept that early targeting of conserved neoantigens can prime broader anti-tumor immunity, a finding of particular relevance for patients with multiple primary tumors or metastatic disease.

Single-cell RNA and TCR sequencing further revealed that this therapeutic strategy not only enhances the recruitment of stem-like and proliferating CD8+ and CD4+ T cells, as observed in other models (*16*, *41*), but also MHC II+ CXCL9+ CXCL10+ CCL5+ CCL8+ TNF+ macrophages (*42*), NK cells, and dendritic cells, while simultaneously reducing immunosuppressive SPP1+ macrophages and exhausted T cells. Vaccination reshaped the T cell repertoire, expanding diverse clonotypes with predicted specificity for conserved neoantigens, and that were distinct from those arising after PD-1 blockade (*42*). Moreover, following vaccination, tumor cells exhibited enhanced expression of ISGs, cytolytic effectors, and antigen presentation genes, alongside suppressed metabolic activity, suggesting that this approach also reprograms the tumor cell transcriptome to favor immune clearance. These ISGs are also highly expressed at baseline in myeloid cells infiltrating patient MMRd tumors (*43*). Altogether, vaccination against conserved neoantigens promoted infiltration of stem-like and proliferating CD4+ and CD8+ T cells, MHC+ macrophages and DCs, while reducing immunosuppressive macrophages and exhausted T cells. PD-1 blockade was characterized by CD8+ T cell, C1Q+ macrophage infiltration, and a diminution of NK cells compared to untreated tumors (*8*). In combination with PD-1 blockade, vaccination enhanced clonal expansion of T cells, induced antigen presentation and interferon-stimulated gene expression in tumors, and synergistically improved anti-tumor immune responses.

Of note, one of the MMRd associated indel mutations that we identified (within the *Igf2r* gene) and predicted candidate neoantigens were found in one of our Lynch CRC patient, and one other (*Chrnb2*) and associated candidate neoantigens were also found in a murine model of Lynch syndrome (*15*). Whole genome sequencing analyses also revealed that untranslated (UTR) and coding regions exhibited the largest fraction of mutations leading to highly immunogenic peptides in *Mlh1* KO murine tumors (*44*). MHC I associated peptides (MAPs) derived from UTRs and out-of-frame translation of coding regions were highly enriched in *Mlh1* KO cells. These MAPs trigger T-cell activation in mice primed with *Mlh1* KO cells (*44*). Thus, our vaccination approaches with shared peptides derived from coding regions may be also relevant in other MMRd contexts such as *Mlh1* epigenetic silencing or Lynch Syndrome, a condition where the risk to develop cancer is increased by 80% (*1*).

Our findings position conserved neoantigens as promising therapeutic targets for pan-tumor immunotherapy in MMRd settings, and suggest a paradigm where vaccination and checkpoint blockade together can amplify adaptive and innate immune responses across tumor types. While our study was conducted in murine models, the strong overlap with human MMRd mutations and neoantigens, including evidence from Lynch Syndrome patients, underscores the translational potential.

In conclusion, our work establishes that conserved neoantigens across organs and species can drive CD8+ T cell-mediated anti-tumor memory responses, enabling abscopal immunity and therapeutic synergy with immune checkpoint blockade. These findings provide a strong rationale for developing conserved neoantigen-targeted vaccines in MMRd tumors and potentially for personalized immunoprevention strategies in high-risk individuals such as those with Lynch Syndrome.

## Acknowledgments

The authors thank all the current and previous members of the Bhardwaj, Samstein, Merad and Vabret labs for critical comments on the manuscript. The authors also thank the members of the Human Immune Monitoring Center for processing the scRNAseq and spatial RNAseq samples and providing the data, the members of the RNA Nanocore, the flow cytometry core, the microscopy core and the pathology core at Mount Sinai Hospital for helpful discussions and training to prepare the experiments. This work was supported in part through the computational and data resources and staff expertise provided by Scientific Computing and Data at the Icahn School of Medicine at Mount Sinai and supported by the Clinical and Translational Science Awards (CTSA) grant UL1TR004419 from the National Center for Advancing Translational Sciences. Research reported in this publication was also supported by the Office of Research Infrastructure of the National Institutes of Health under award number S10OD026880 and S10OD030463. The content is solely the responsibility of the authors and does not necessarily represent the official views of the National Institutes of Health. We especially thank Tim O’Donnel and Mesude Bicak for discussions about MHC flurry, Flora Borne for discussions about WES and bulkRNAseq, and Ashley Reid Cahn for discussions about the Elispot protocol.

## Author Contributions

G.M. and N.B. conceptualized the project. G.M. performed the in vivo and in vitro experiments. G.M. performed the analysis of the data. R.W.W helped with the production of vaccine doses. N.Vaninov helped with the in vivo orthotopic experiments for the 4T1 model. J.B. and F.R. helped with the in vivo orthotopic experiments for the CT26 model. M.B. and A.L. helped to identify one patient with the mutation of interest in Mount Sinai Lynch cohort. E.O and M.S. provided the C11 cell line and P.S. provided GFP tetramers. S.B. sorted GFP tumor cells. L.V. and A.A. provided multiple reagents for the experiments. Z.C. helped for the processing of scRNAseq samples. C.C.B. and N.Vabret helped with the vaccine design. G.M. wrote the manuscript. G.M., C.C.B. and N.B. revised the manuscript.

## Methods

### Human subjects protocol overview and patient information

Patients from the Mount Sinai Health System with evidence of a germline mutation associated with Lynch syndrome (MLH1, MSH2, MSH6, PMS2, and EPCAM) or the development of MSI-H cancer (with immunohistochemical verification of MMR protein loss) undergoing treatment or routine surveillance screening (along with MSS controls) were recruited to our active IRB protocols (21-01317 and 19-00936).

### MSH2-deficient tumor cells

CT26 and 4T1 cells deleted for MSH2 were provided by R.Samstein (*7*). Guide RNA sequences (5’-CGGTGCGCCTCTTCGACCGC-3’) and (5’- GCACGGAGAGGACGCGCTGC-3’) targeting mouse Msh2 exon 1 were cloned into the PX461 plasmid and co-transfected into mouse CT26 mouse colon carcinoma cells using GenJetTM (SignaGen®) In Vitro DNA Transfection Reagent following the manufacturer’s protocol. 48 hours following transfection, GFP+ cells were seeded at one cell per 96-well by the Flow Cytometry Core Facility (FCCF). Cells were grown in RPMI supplemented with 10% FBS. Single cell clones were expanded and depletion of MSH2 was confirmed by Western Blot (anti-MSH2 monoclonal antibody (FE11), Invitrogen Antibodies). Confirmed Msh2-/- single cell subclones were expanded and used for serial passaging and downstream in vivo studies further described below.

### Cell culture

Cells were amplified in RPMI media (Sigma) or DMEM media (Gibco) for cell culture. 10% FBS, 2 mM L-glutamine (Gibco) and 10% gentamicin were added to the medium. Medium was renewed 2 times a week. For cell amplification, cancer cells were seeded at 1,000 cells/cm2 and sub-cultured every week. All cultures were performed in plastic flasks (Biocoat, Becton-Dickinson). CT26 cells were serially passaged for continuous lengths of time under standard tissue culture conditions and were frozen under standard cell-specific freezing conditions with heat inactivated FBS and 10% DMSO cryoprotectant. Depletion of MSH2 was confirmed by Western Blot (anti-MSH2 monoclonal antibody (FE11), Invitrogen Antibodies). C11 is a primary lung tumor from a male BALB/c mouse 77 weeks after urethane injection was minced and passaged through an NSG mouse for 2 months. The resulting tumor was then digested and a cell line was established after 6 passages in culture.

### In vivo tumor growth analysis

2 * 10^5 WT CT26 or 4T1 cells or CT26 or 4T1 MSH2 KO cells in 100 µl of PBS were injected into the flanks of 6-week old BALB/c mice (Jackson Laboratories). For orthotopic injections, these tumor cells were injected in fat pad (4T1) or in 10 μL of matrigel (BME) in the colon (CT26). Tumor volumes were measured weekly and calculated by the formula: (1/2) * (𝐿𝑒𝑛𝑔𝑡ℎ) * (𝑊𝑖𝑑𝑡ℎ)² . 5 to 20 mice were used for each group. Mice were euthanized by carbon dioxide and the flank tumors were resected 21 days after tumor injection.

### Spheroid growth

10,000 tumor cells were seeded in 200µL of RPMI medium in a nucleon sphera 96 well plate (Thermofisher). As an alternative, spheroids were grown using SpheroTribe (Idylle) (*45*). After 1 to 3 days, the circularity and the area of spheroids were measured as previously described in other systems (*46*). Circularity = (4π Area) / (perimeter)², Roundness = 4 Area / (𝜋 × major axis²), with Area in µm². All parameters were measured using the Image J software.

### Spheroid infiltration and killing by PBMC

10,000 cells were seeded in 96 low attachment well plates (Nucleonsphera, Thermofisher) with RPMI medium, to form spheroids. PBMC were seeded in other 96-well culture plates (Thermofisher) and incubated at 37 °C, in 5% CO2 in 200 μL RPMI medium supplemented with 20% FCS, enriched in gentamicin. 100,000 PBMCs were activated per well by IL-15 (40ng/mL) for 3 days. Then, these 100,000 PBMC were added to each tumor spheroid (10,000 cells) for 3 days, in presence or not with peptides (10nM). Spheroids incubated with PBMC were isolated from PBMCs in suspension and dissociated using MACS dissociator and a mouse tumor dissociation kit (Miltenyi). Then, PBMC, spheroids or spheroids incubated with activated PBMC were labeled with antibodies listed in **Table 0** and Live Dead Blue (1uL of each antibody in 200uL of PBS). The collection was made using an Attune NxT flow cytometer (Thermofisher) or a spectral Aurora (Cytek). Data were analyzed using FlowJo software (BD biosciences). PBMC ability to infiltrate spheroids was analyzed by comparing CD45 expression on spheroids incubated with PBMC in different conditions. Tumor cell death was measured using propidium iodide (5μg/mL) as previously described (*46*).

### Flow cytometry analysis

Fresh tumors were resected and dissociated into single cell suspensions using a gentle MACS tissue dissociator and mouse tumor dissociation kit (Miltenyi). For analysis of cell-surface immune markers expression, cancer cells were processed as single-cell suspensions and stained for 30 minutes at room temperature with monoclonal antibodies. PBMC were labeled with antibodies listed in **Table 0** and Live Dead Blue (1uL of each antibody in 200uL of PBS). Non-reactive antibodies of similar species and isotype, and coupled with same fluorochromes, were used as isotypic controls. Immune expression profiles were analyzed using an Attune NxT (Thermofisher) or a spectral Aurora (Cytek). Data were analyzed using FlowJo software (BD biosciences).

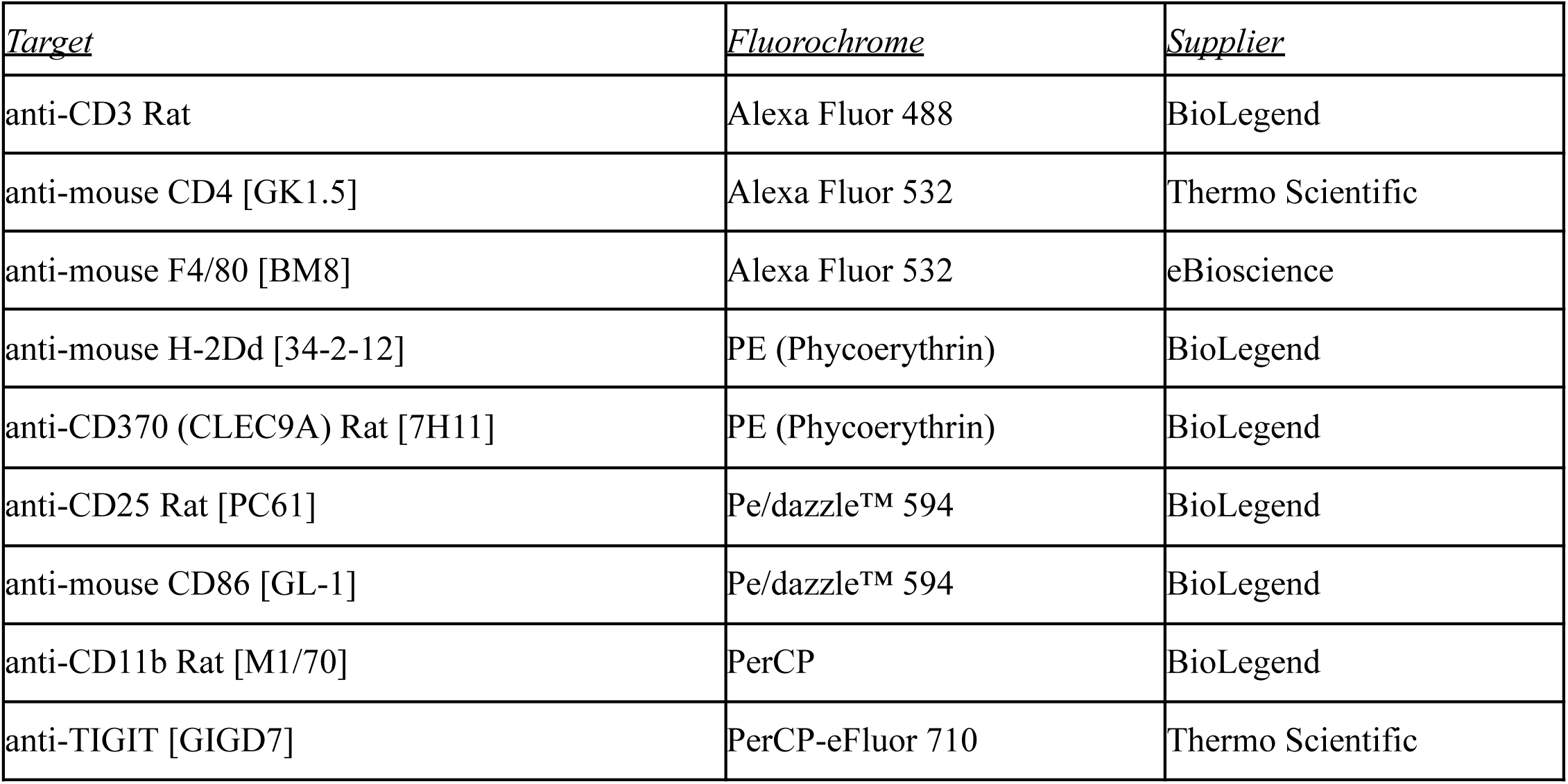
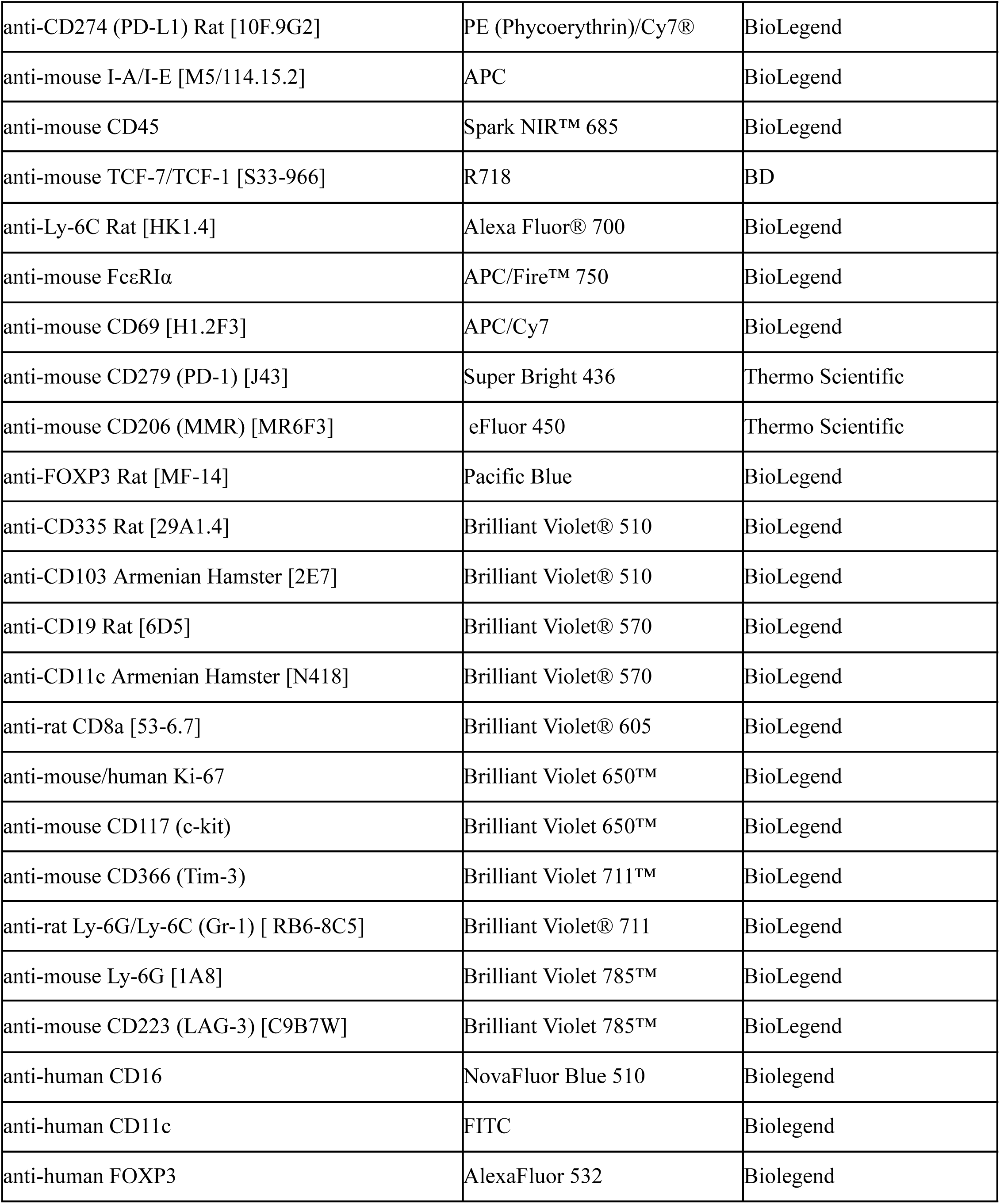
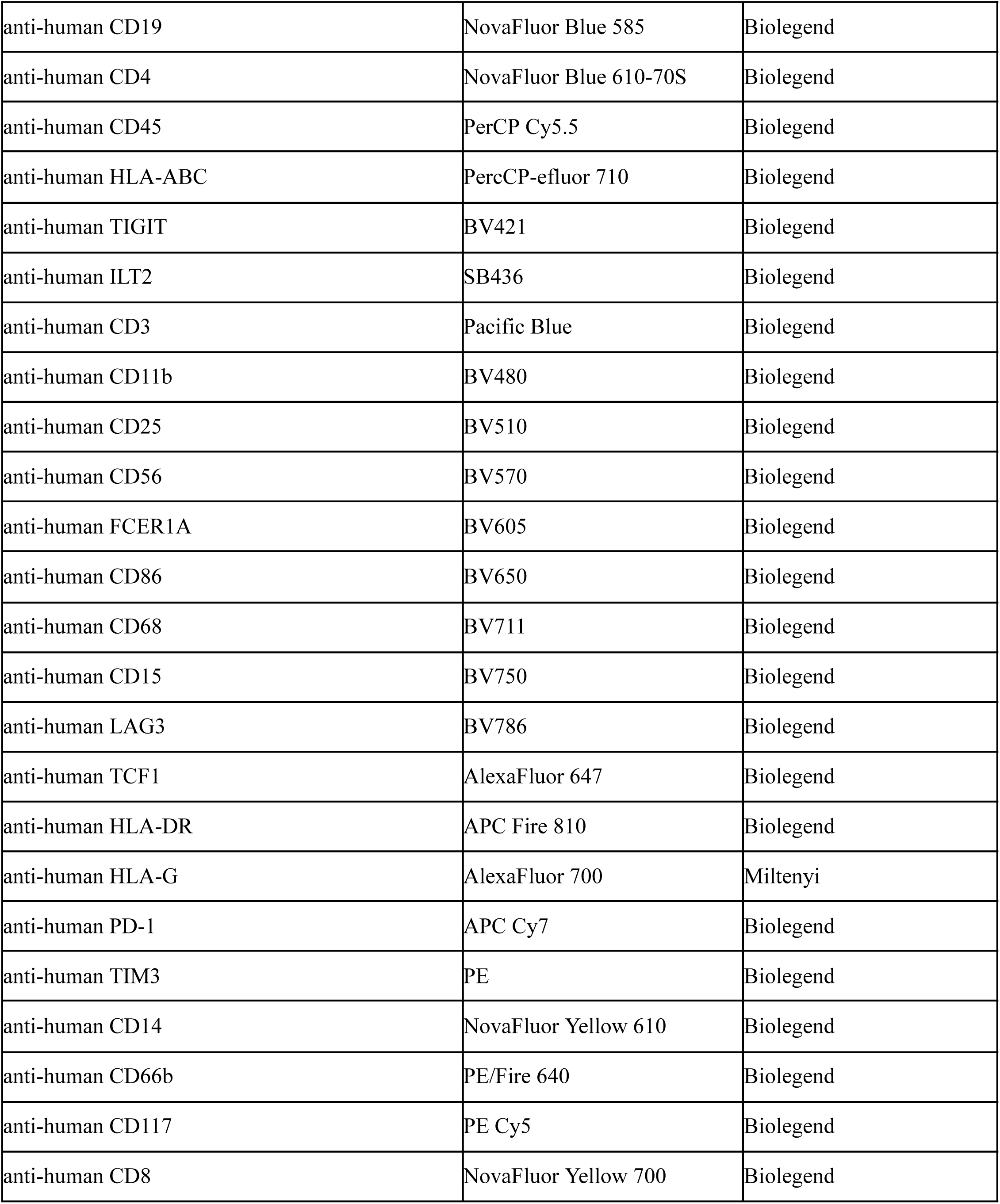
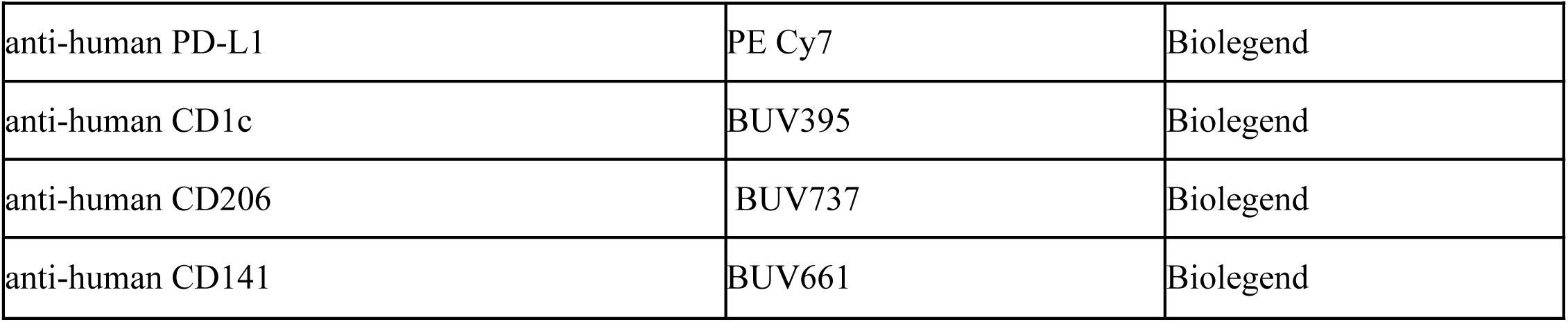

| **<u>Table 0) List of antibodies.</u>** |  |  |
| --- | --- | --- |
| *<u>Target</u>* | *<u>Fluorochrome</u>* | *<u>Supplier</u>* |
| anti-CD3 Rat | Alexa Fluor 488 | BioLegend |
| anti-mouse CD4 [GK1.5] | Alexa Fluor 532 | Thermo Scientific |
| anti-mouse F4/80 [BM8] | Alexa Fluor 532 | eBioscience |
| anti-mouse H-2Dd [34-2-12] | PE (Phycoerythrin) | BioLegend |
| anti-CD370 (CLEC9A) Rat [7H11] | PE (Phycoerythrin) | BioLegend |
| anti-CD25 Rat [PC61] | Pe/dazzle™ 594 | BioLegend |
| anti-mouse CD86 [GL-1] | Pe/dazzle™ 594 | BioLegend |
| anti-CD11b Rat [M1/70] | PerCP | BioLegend |
| anti-TIGIT [GIGD7] | PerCP-eFluor 710 | Thermo Scientific |
| **Table 0) List of antibodies.** |  |  |
| anti-CD274 (PD-L1) Rat [10F.9G2] | PE (Phycoerythrin)/Cy7® | BioLegend |
| anti-mouse I-A/I-E [M5/114.15.2] | APC | BioLegend |
| anti-mouse CD45 | Spark NIR™ 685 | BioLegend |
| anti-mouse TCF-7/TCF-1 [S33-966] | R718 | BD |
| anti-Ly-6C Rat [HK1.4] | Alexa Fluor® 700 | BioLegend |
| anti-mouse FcεRIα | APC/Fire™ 750 | BioLegend |
| anti-mouse CD69 [H1.2F3] | APC/Cy7 | BioLegend |
| anti-mouse CD279 (PD-1) [J43] | Super Bright 436 | Thermo Scientific |
| anti-mouse CD206 (MMR) [MR6F3] | eFluor 450 | Thermo Scientific |
| anti-FOXP3 Rat [MF-14] | Pacific Blue | BioLegend |
| anti-CD335 Rat [29A1.4] | Brilliant Violet® 510 | BioLegend |
| anti-CD103 Armenian Hamster [2E7] | Brilliant Violet® 510 | BioLegend |
| anti-CD19 Rat [6D5] | Brilliant Violet® 570 | BioLegend |
| anti-CD11c Armenian Hamster [N418] | Brilliant Violet® 570 | BioLegend |
| anti-rat CD8a [53-6.7] | Brilliant Violet® 605 | BioLegend |
| anti-mouse/human Ki-67 | Brilliant Violet 650™ | BioLegend |
| anti-mouse CD117 (c-kit) | Brilliant Violet 650™ | BioLegend |
| anti-mouse CD366 (Tim-3) | Brilliant Violet 711™ | BioLegend |
| anti-rat Ly-6G/Ly-6C (Gr-1) [RB6-8C5] | Brilliant Violet® 711 | BioLegend |
| anti-mouse Ly-6G [1A8] | Brilliant Violet 785™ | BioLegend |
| anti-mouse CD223 (LAG-3) [C9B7W] | Brilliant Violet 785™ | BioLegend |
| anti-human CD16 | NovaFluor Blue 510 | Biolegend |
| anti-human CD11c | FITC | Biolegend |
| anti-human FOXP3 | AlexaFluor 532 | Biolegend |
| **Table 0) List of antibodies.** |  |  |
| anti-human CD19 | NovaFluor Blue 585 | Biolegend |
| anti-human CD4 | NovaFluor Blue 610-70S | Biolegend |
| anti-human CD45 | PerCP Cy5.5 | Biolegend |
| anti-human HLA-ABC | PercCP-eFluor 710 | Biolegend |
| anti-human TIGIT | BV421 | Biolegend |
| anti-human ILT2 | SB436 | Biolegend |
| anti-human CD3 | Pacific Blue | Biolegend |
| anti-human CD11b | BV480 | Biolegend |
| anti-human CD25 | BV510 | Biolegend |
| anti-human CD56 | BV570 | Biolegend |
| anti-human FCER1A | BV605 | Biolegend |
| anti-human CD86 | BV650 | Biolegend |
| anti-human CD68 | BV711 | Biolegend |
| anti-human CD15 | BV750 | Biolegend |
| anti-human LAG3 | BV786 | Biolegend |
| anti-human TCF1 | AlexaFluor 647 | Biolegend |
| anti-human HLA-DR | APC Fire 810 | Biolegend |
| anti-human HLA-G | AlexaFluor 700 | Miltenyi |
| anti-human PD-1 | APC Cy7 | Biolegend |
| anti-human TIM3 | PE | Biolegend |
| anti-human CD14 | NovaFluor Yellow 610 | Biolegend |
| anti-human CD66b | PE/Fire 640 | Biolegend |
| anti-human CD117 | PE Cy5 | Biolegend |
| anti-human CD8 | NovaFluor Yellow 700 | Biolegend |

| **Table 0) List of antibodies.** |  |  |
| --- | --- | --- |
| anti-human PD-L1 | PE Cy7 | Biolegend |
| anti-human CD1c | BUV395 | Biolegend |
| anti-human CD206 | BUV737 | Biolegend |
| anti-human CD141 | BUV661 | Biolegend |

### Bi-photon spheroid imaging

CT26 spheroids deleted for MSH2 were grown for 3 days and then co-cultured with or without immune cells for 3 more days. Blood immune cells are previously labeled by Life Technologies celltracker Deep Red (Thermo Scientific). Spheroid cell nuclei are labeled with Molecular Probes NucBlue Fixed Cell ReadyProbes Reagent (Thermo Scientific). 3D Bi-photon imaging of spheroid infiltration by immune cells was performed using Olympus FVMPE-RS. Spheroid immune infiltration was followed on the (x/y) axis in the z-middle. Z-stack imaging of spheroid immune infiltration (top to middle of the spheroid) was also performed.

### Single cell mRNA and TCR V(D)J sequencing

CT26 WT and MSH2 KO tumors were inoculated in BALC mice for 4 weeks as described above. Tumors were harvested after 4 weeks and dissociated into single cell suspensions.12 tumors (3 per condition) were used for single-cell RNA sequencing performed at the Human Immune Monitoring Center (HIMC) at Mount Sinai. Viability of single cells was assessed using Acridine Orange/Propidium Iodide viability staining reagent (Nexcelom), and debris-free suspensions of >75% viability were deemed suitable for the experiments. Tumor cells from individual mice were labeled with unique TotalSeqC hashtag antibodies (BioLegend) and pooled in equal proportion. scRNAseq was performed using the Chromium platform (10x Genomics) with the 5’ gene expression (5’ GEX) V2 kit, with a targeted recovery of 10,000 cells/lane. Each pool was loaded on three lanes to increase total cell yield. Gel-Beads in Emulsions (GEMs) were generated on the sample chip in the Chromium X system. Barcoded cDNA was extracted from the GEMs after Post-GEM RT-cleanup and amplified for 13 cycles. For GEX libraries, amplified cDNA was fragmented and subjected to end-repair, poly A-tailing, adapter ligation, and 10x-specific sample indexing following the manufacturer’s protocol. For TCR V(D)J libraries, the cDNA was used as a template and the specific variable regions (V, D, and J regions) from each TCR locus are amplified and indexed following the manufacturer’s protocol. Hashtab library was prepared per manufacturer’s instructions (10x Genomics). All libraries were quantified using TapeStation (Agilent) and QuBit (ThermoFisher) analyses and were sequenced in paired end mode on a NovaSeq instrument (Illumina) targeting a depth of 20,000 reads per cell for GEX and 5,000 reads per cell for TCR and Hashtags. Raw fastq files were aligned to the mm10 reference genome (2020-A) reference genome and demultiplexed using Cell Ranger multi v7.0.1 (10x Genomics). Downstream analysis of 55,814 cells was performed using python and scanpy. All samples were preprocessed to remove genes that are expressed in less than 3 cells and to identify mitochondrial genes. Cells that have >20 mitochondrial genes were removed from the analysis. Data were normalized and highly-variable genes were identified. Principal component analysis and clustering were performed using the leiden graph-clustering method. Marker genes were identified for each cluster and a secondary clustering was performed to separate the immune cells and the cancer cells with the Wilcoxon

Rank-Sum test. Gene expression was analyzed in the different conditions using scanpy. Single cell clustering was finally applied to the spatial samples using scanorama and scanpy.

### Visium Spatial Gene Expression analysis

CT26 WT and MSH2 KO tumors were inoculated for 4 weeks in BALC mice as described above. Tumors were harvested after 4 weeks, and 4 tumors (2 per condition) were fixed in 4% PFA and incorporated in paraffin blocks. RNA quality was assessed using TapeStation (Agilent) and confirmed to be suitable for the assay with DV200 values > 30%. Spatial Gene Expression assay was performed using Visium CytAssist platform following manufacturer’s instructions (10x Genomics. Briefly, 5 um FFPE tissue sections that were placed on a plain glass slide were deparaffinized and decrosslinked. The mouse whole transcriptome probe panel consisting of specific probes for each targeted gene was then added to the tissue. These probe pairs hybridize to their gene target and are then ligated to one another. Tissue slides and Visium CytAssist Spatial Gene Expression Slides were loaded into the Visium CytAssist instrument, where they were brought into proximity with one another. Each Visium CytAssist Slide contains about 5,000 Capture Areas (55 um across) with barcoded spots that include oligonucleotides required to capture gene expression probes. Gene expression probes were released from the tissue upon CytAssist Enabled RNA Digestion & Tissue Removal, enabling capture by the spatially barcoded oligonucleotides present on the Visium slide surface. All the probes captured on a specific spot share a common Spatial Barcode. Libraries were generated from the probes and sequenced, and the Spatial Barcodes were used to associate the reads back to the tissue section images for spatial mapping of gene expression. Sequencing was performed on an Illumina NovaSeq sequencer with depth set at 25,000 read per capture spot. Fastq files were subsequently loaded on to Space Ranger (10x Genomics) for read alignment, spot detection, gene expression quantification and data visualization. SpatialRNAseq data were analyzed using Squidpy, Scanorama and Scanpy.

### Whole Exome Sequencing (WES)

A sample sheet was generated to document murine sample details, including sample paths, using a format specifying murine ID, sample type (normal or tumor), lane information, and corresponding FASTQ files. The sample sheet, named "samplesheet.csv," was organized within the directory containing the associated FASTQ files. A job file, denoted as "submit_job.sh," was created to facilitate variant calling using Nextflow. This job file incorporated necessary module additions for Java (version 11.0.2) and Singularity (version 3.2.1). The Nextflow script executed the variant calling pipeline from nf-core/sarek, tailored for WES data analysis (*47*, *48*). Input parameters included the path to the sample sheet (--input), maximum CPU allocation (--max_cpus), specification for only paired variant calling (--only_paired_variant_calling), selected variant calling tools (e.g., Strelka, Mutect2), reference genome (e.g., GRCm38), and output directory (--outdir). Nextflow was set up by installing the required modules for Java (version 11.0.2) and Singularity (version 3.2.1). The Nextflow executable was downloaded and configured in the user’s home directory (∼/nextflow). Verification of the Nextflow installation and version was performed to ensure proper functionality. Upon setup completion, variant calling jobs were initiated for each sample using the provided job file (submit_job.sh). These jobs were submitted to the computing cluster for execution, with subsequent monitoring until completion. Variant annotation was conducted using Python scripting with the Varcode library. Variant Call Format (VCF) files were loaded, correcting genomic mismatches as necessary. Non-protein-modifying mutations were filtered out, and mutations were annotated with corresponding gene information. Original and mutant protein sequences were retrieved, followed by segmentation into 9-mers. Frameshift mutations were identified and mutant-specific 9-mers were listed.

### Bulk RNA-Seq to verify the expression of mutated transcripts

To verify the expression of somatic mutations at the transcript level, bulk RNA sequencing (RNA-seq) was performed on tumor tissue samples. Cell pellets were sent to Azenta for sequencing following total RNA extraction libraries preparation (paired-end reads). Raw reads were quality-checked with FastQC and trimmed using Trim_galore. Alignment to the murine reference genome (GRCm39) was performed using STAR. Somatic mutations previously identified by matched WES were used as a reference list. BAM files were inspected for variant-supporting reads at corresponding genomic loci using Samtools and IGV. Only mutations with ≥3 uniquely mapped variant-supporting reads were considered as transcriptionally expressed.

### Alpha Fold prediction of protein structure according to mutation status

Protein structures were predicted using Alpha Fold algorithms (*49*) and the amino acid sequences of the protein of interest. We showed the structure of the protein with the highest probability. The number of sequences per position is superior to 30 sequences per position. The predicted lDDT per position represents the model confidence (out of 100) at each position. We used the PyMol software to show the structures of the mutated and WT proteins aligned together.

### Peptide selection

Peptide prediction was performed using the MHCflurry Colab environment (*50*, *51*). The provided code was customized with mutated peptides and specified murine mhc alleles. Peptide affinities for MHC binding were predicted accordingly. Predicted peptides were filtered based on predefined criteria, including MHC binding affinity (<500nM), shared presence across different tumors, low presentation percentile (<2%), high presentation score, and procession score.

### mRNA vaccines

Plasmids containing the shared FS indel mutations of interest were generated as previously described (*30*, *31*). For each mutation, we selected the sequence starting with 8 amino acids before the indel mutation and all the downstream mutated sequence to maximize the probability to generate immunogenic 9mers and 15mers. Each sequence was separated by a glycine-serine (GS) linker sequence, and we added a model epitope derived from ovalbumin (OVA). The mRNA sequences also include a 5′ untranslated region (UTR), a 3′UTR and a polyadenylation (PolyA) tail. An adenine (A) nucleotide was engineered upstream of the 5′UTR to enable effective capping of the in vitro transcribed mRNAs using the CleanCap AG (TriLink BioTechnologies). The plasmid DNA encoding the mRNA templates were amplified with E.coli transformation and linearized with BsPQI (New England Biolabs). Linearization was verified by electrophoresis with initial plasmids and linearized DNA. Corresponding mRNA, IVT-mRNA lipid nanoparticles (LNPs) were produced as previously described (*31*), and characterization of antigen presentation after transfection of mRNA vaccine into the B16F10 cancer cell line was performed using OVA (*30*). In prophylactic studies, we immunized each mouse with 5 μg of mRNA-LNPs diluted in 50 μL 1x PBS via retro-orbital injection at day 0. For negative control, immunize mice with vehicle control (1X PBS). A booster dose was given on day 5 with 5 μg mRNA-LNPs or PBS as negative control. Then, mice were challenged with tumors on day 10. In therapeutic studies, mice were challenged with tumors on day 0, and we immunized each mouse with 5 μg of mRNA-LNPs diluted in 50 μL 1x PBS via retro-orbital injection at day 5. For negative control, we immunized mice with a vehicle control (1X PBS). A booster dose was given on day 10 with 5 μg mRNA-LNPs or PBS as negative control. IgG or ICB antibodies such as anti-mPD-1 Invivomab anti-mouse (CD279) (Bio X Cell) (100µg) were administered intraperitoneally in 100 µl of PBS every 3-4 days, starting after day 7.

### Elispot assay

We performed an enzyme-linked immunospot (ELISPOT) assay to quantify antigen-specific T cell responses. Ninety-six-well plates were coated overnight with anti-IFN-γ antibodies and blocked with RPMI 1640 containing 10% fetal bovine serum. Blood cells from vaccinated mice were isolated and resuspended in culture media at a concentration of 10^7 cells/mL. Each well received 100 µL of cell suspension and specific peptides identified in Table 1 or the WT peptides (2.5µg/mL), with controls including aCD28 and aCD49d co-stim only and mitogen-stimulated wells (PMA/iono). After incubating for 24 hours at 37°C with 5% CO₂, cell suspensions were removed, and plates were washed. Detection involved adding biotinylated secondary antibodies followed by streptavidin-HRP, with substrate added to visualize spots. Plates were analyzed for spot-forming units (SFUs) to assess cytokine secretion.

### Neoantigen-specific T cell expansion and functional characterization

T cell expansion assays were utilized to characterize neoantigen peptide immunogenicity and polyfunctional cytokine responses elicited by CD8+ and CD4+ T cell subsets as described previously (*52*). Prior to culture, cryopreserved PBMCs were thawed in 37°C water bath and transferred into RPMI medium (11875-093, Gibco) containing DNase I (Sigma-Aldrich) at a final concentration of 2 U/mL, centrifuged at 500 g for 5 min and resuspended in RPMI. Then PBMCs were resuspended in X-VIVO 15 medium (02-053Q, Lonza), seeded at a density of 10^5 cells per well in U-bottom 96-well plates and cultured with cytokines promoting differentiation of dendritic cell (DC) subsets: GM-CSF (0024-5843-05, SANOFI, 1000 IU/mL), IL-4 (204-IL-010, R&D Systems, 500 IU/mL) and FLT3L (308-FKN-025, R&D Systems, 50 ng/mL). PBMCs were cultured for 24 hours before stimulation with control or neoantigen peptides (Extended Figure 2 for full peptide sequences) at a final concentration of 2.5 μM in combination with DC maturation adjuvants: LPS (201-LB-010, Invivogen, 0.1 mg/mL), R848 (tlrl-r848-5, Invivogen, 10 mM) and IL-1β (201-LB-010, R&D Systems 10 ng/mL), in X-VIVO 15 medium. The cells were given (every 2-3 days beginning 24 hours after the initial stimulation) cytokines promoting T cell expansion, IL-2 (202-IL-050, R&D Systems, 10 IU/mL), IL-7 (207-IL-025, R&D Systems, 10 ng/mL) and IL-15 (200-15, Peprotech, 10 ng/mL) in complete RPMI media (GIBCO) containing 10% human serum (R10). Cells were harvested, wells were pooled within expansion groups and then cell were washed before resuspension in R10 following 10 days of culture. Expanded cells were then seeded at 2 x10^5 cells per well in U-bottom 96-well plates and re-stimulated with 2.5 μM control or neoantigen peptides together with 0.5 mg/mL of costimulatory antibodies, anti-CD28 (555726, BD Biosciences) and anti-CD49d (555502, BD Biosciences), and protein transport inhibitors BD GolgiStopTM (554724), containing monensin and BD GolgiPlugTM (555029), containing brefeldin A according to manufacturer’s instructions. Cells were processed for intracellular staining for flow cytometry using BD Cytofix/CytopermTM reagents according to manufacturer’s protocol. The following antibodies were used: for surface staining CD3 (clone SK7, FITC), CD4 (clone RPA-T4, BV785) and CD8a (clone RPA-T8, APC) and for intracellular staining IFN-g (clone B27, PE), and TNF-a (clone Mab11, PE/Cy7). All antibodies were purchased via BioLegend. LIVE/DEAD Fixable Blue Dead Cell Stain Kit by Thermo Scientific was used for live and dead cell discrimination. Data was acquired using the Attune. FlowJo V10 was used for analysis. DMSO (Sigma-Aldrich) was used at the equal volume of the test peptides and served as the vehicle/negative control (at least 2 replicates per test group).

### Data analysis and visualization

Spectral flow cytometry and RNAseq data were analyzed using Python 3 and Jupyter Notebook. Numpy and Pandas were used for array and data frame operations, and data visualization was performed using Matplotlib and Seaborn. scRNAseq and spatialRNAseq data were analyzed using Cell Ranger, Squidpy, Scanorama and Scanpy. Flow cytometry data were analyzed using FlowJo and Python 3.

### Statistics

Statistical significance of the observed differences was determined using the Mann-Whitney two-sided U-test, T-test or Linear regression. All data are presented as mean±SEM. The difference was considered as significant when the p value was below 0.05 *.

## Resource Availability

### Lead Contact

Further information and requests for the resources and reagents should be directed to and will be fulfilled by the lead contact, Nina Bhardwaj.

### Materials Availability

This study did not generate new unique reagents.

### Data and Code Availability

Data reported in this study are tabulated in the main text and supplementary materials. Code is available on Github and GoogleColab.

https://github.com/gmestrallet/BasicScRNAseq

https://github.com/gmestrallet/BasicSpatialRNAseq https://github.com/gmestrallet/TCRscRNAseqAnalysis

## Extended data

**Extended Figure 1.**
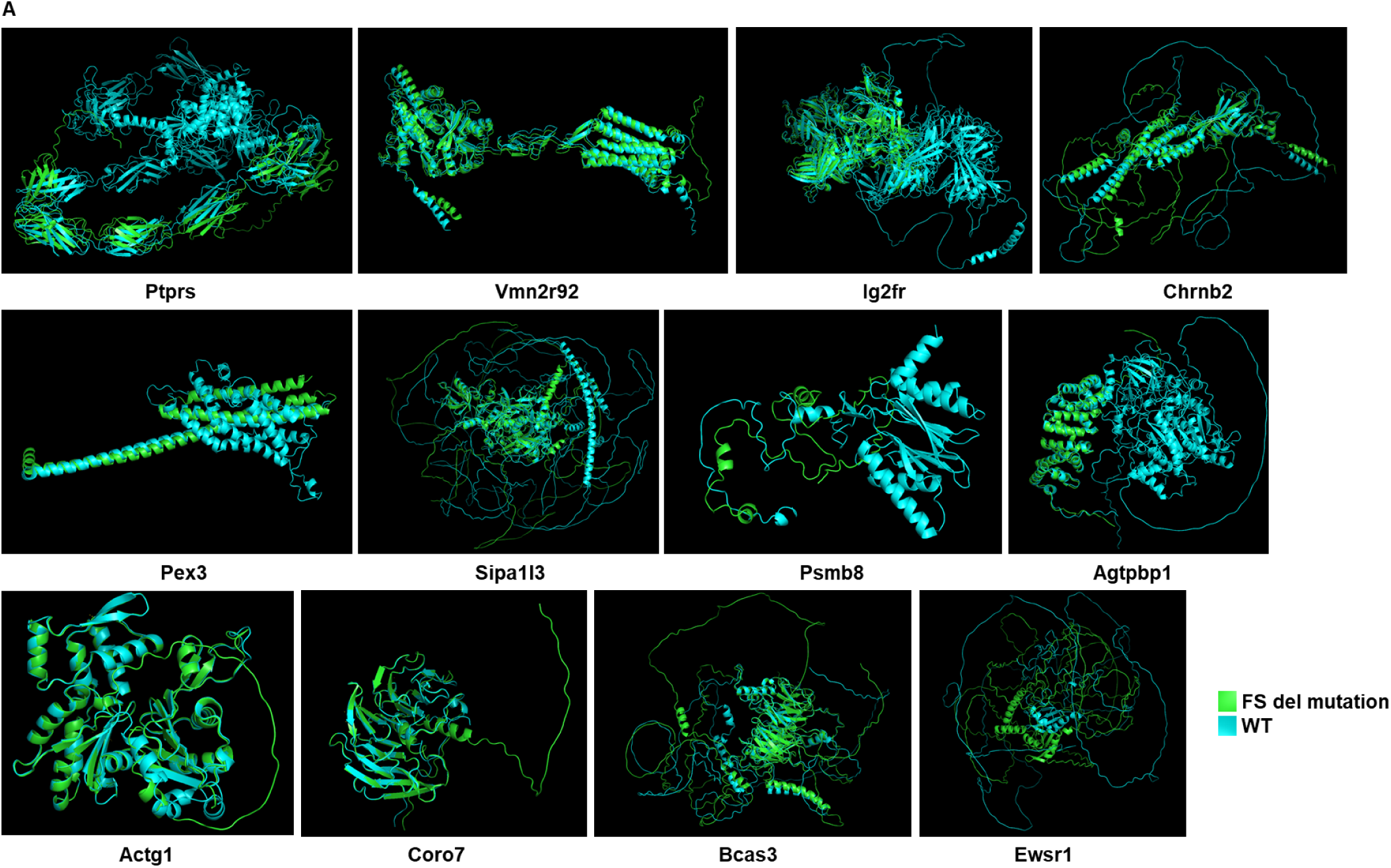
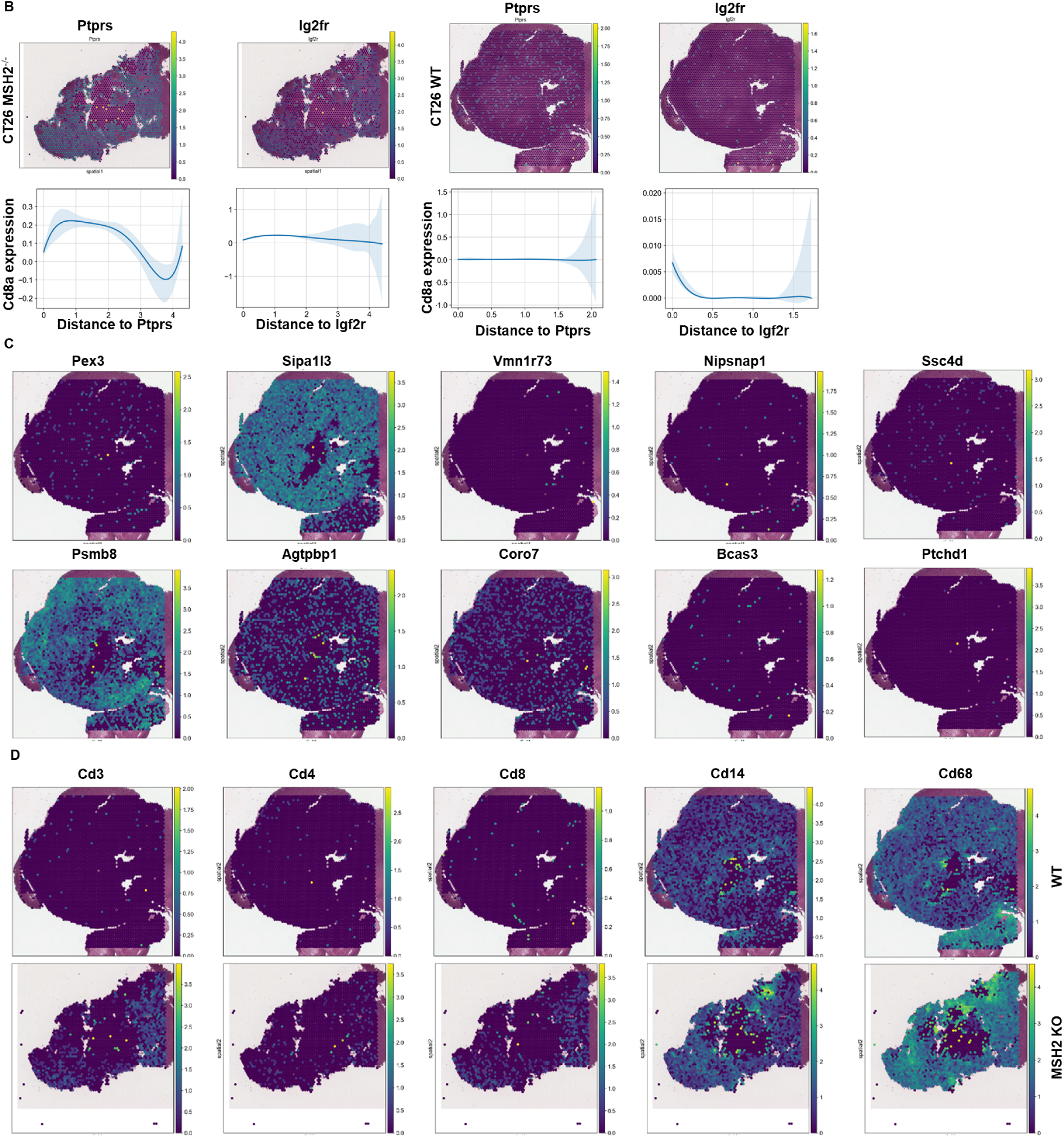
Protein structure properties and spatial gene expression of the candidate neoantigens and colocalization with immune genes. **A** Prediction of the protein structures according to the presence of FS del mutation of interest using Alphafold. **B** SpatialRNAseq, expression of the genes encoding candidate MSH2 KO neoantigens in CT26 MSH2 KO and WT tumors and colocalization with Cd8 expression. **C** SpatialRNAseq, expression of the genes encoding candidate CT26 neoantigens in a CT26 WT tumor. **D** SpatialRNAseq, expression of the immune genes in a CT26 MSH2 KO tumor and a CT26 WT tumor.

**Extended Figure 2.**
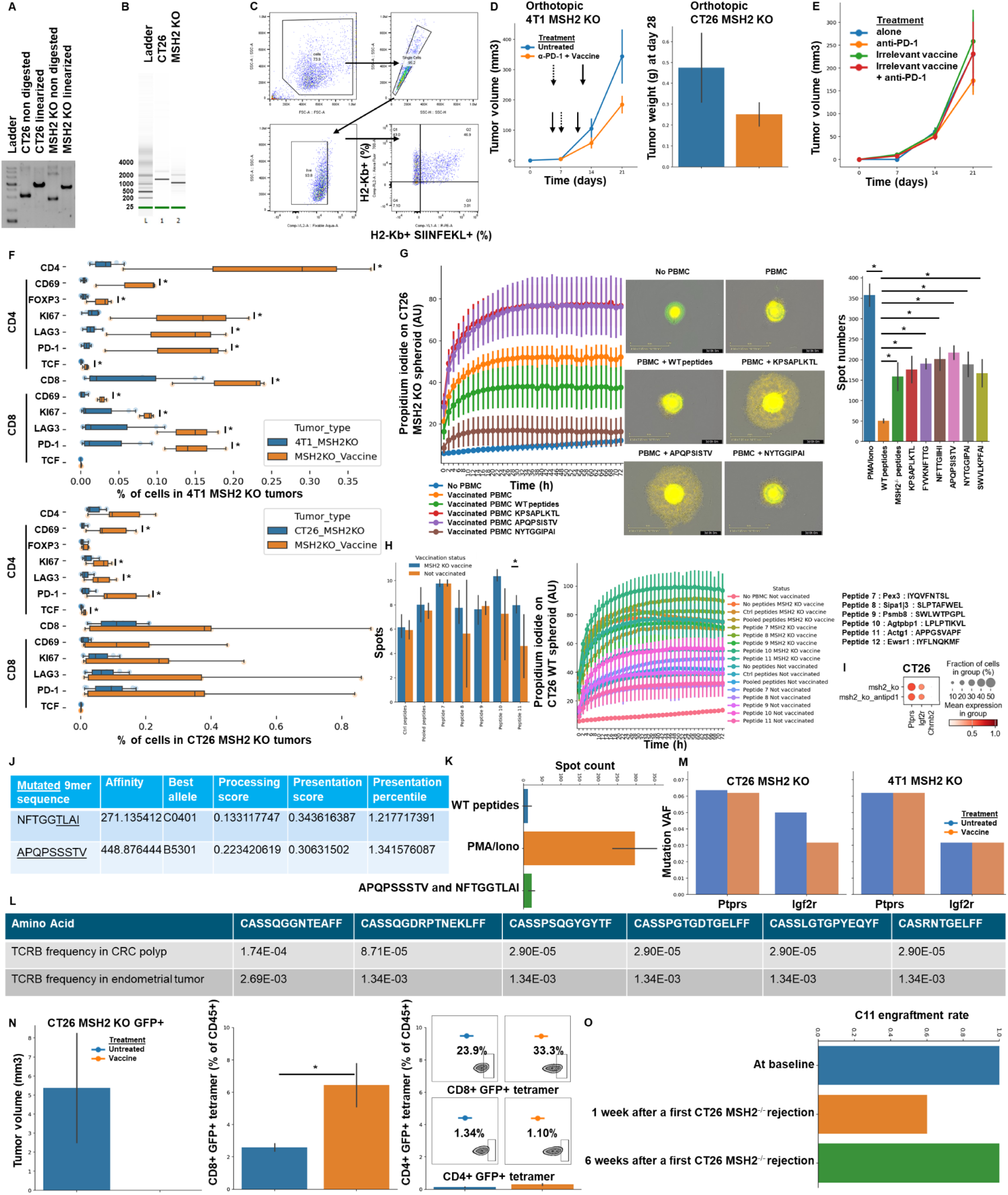
Characterization of mRNA vaccine and shared peptides immunogenicity. **A** Gel electrophoresis of linearized plasmid compared to non-digested plasmid (ND). **B** Bioanalyzer electrophoresis mRNA. **C** B16F10 antigen presentation assay following stimulation with MSH2 KO RNA-Lipoplexes formulations. Flow cytometry gating strategy to quantify antigen presenting (H2-Kb/SIINFEKL+) B16F10 cells. **D** Tumor volume or weight of CT26 MSH2 KO or 4T1 MSH2 KO orthotopic tumors according to the presence of anti-PD-1 (100 ug twice a week 7 days after tumor inoculation) and mRNA vaccine encoding shared MMRd neoantigens (5 μg mRNA-LNPs at 5 days and 10 days after tumor inoculation). N=5 to 6, Mann-Whitney U Test, pvalue<0.05. Data are represented as mean ± SEM. **E** 4T1 MSH2 KO tumor growth following vaccination with mRNA/LNPs vaccine encoding neoantigens not present in 4T1 tumors. **F** Infiltration of 4T1 and CT26 MSH2 KO tumors by immune cells on mice with prophylactic vaccination (flow cytometry). N=3 to 8, Mann-Whitney U Test, pvalue<0.05. Data are represented as mean ± SEM. **G** Spheroid killing and Elispot assays were performed with blood cells from vaccinated mice and stimulation with WT or mutated peptides shared by MSH2 KO tumors and predicted to be immunogenic identified in **Table 1**. N=4, Mann-Whitney U Test, pvalue<0.05. 10,000 CT26 MSH2 KO cells (GFP+) per spheroid, cocultured with 100,000 blood cells from vaccinated mice and stimulated with IL-15 and peptides. Death is measured with propidium iodide. N=4, Mann-Whitney U Test, pvalue<0.05. **H** Elispot and CT26 WT spheroid killing assays were performed with blood cells from vaccinated mice and stimulation with WT or mutated peptides shared by CT26 tumors regardless MSH2 status and predicted to be immunogenic identified in **Table 2**. N=4, Mann-Whitney U Test, pvalue<0.05. **I** ScRNAseq expression of genes encoding candidate MSH2 KO neoantigens identified in table 1 by WES and MHCflurry. **J** Neoantigens from *Ptprs* P622LFs and *Igf2r* D1317TFs mutations shared in our human Lynch and MSI-H tumor cohort, with predicted affinity, best allele binding, processing score, presentation score and percentile by MHCflurry. Igf2r D1317TFs mutation is expressed in the CRC tumor of one of our Lynch patients (HLA-type: A*23:01 A*36:01 B*57:01 B*53:01 C*07:01 C*04:01). **K** Elispot was performed with blood from this Lynch CRC patient and stimulation with WT or mutated peptides shared by MSH2 KO tumors and predicted to be immunogenic. N=4, Mann-Whitney U Test, pvalue<0.05. **L** TCRs shared by several primary tumors on the same patient. **M** VAF of mutated *Ptprs* and *Igf2r* before and after vaccination. **N** CT26 MSH2 KO cells were transduced to express GFP. Tumor volume was measured after 3 weeks according to therapeutic vaccination targeting MMRd neoantigens. Flow cytometry was performed for GFP-tetramer-H2-Kd staining in the spleen of these mice. N=4, Mann-Whitney U Test, pvalue<0.05. Data are represented as mean ± SEM. **O** C11 tumor engraftment rate at baseline, 1 week or 6 weeks after a first CT26 MSH2 KO rejection. N=5.

**Extended Figure 3.**
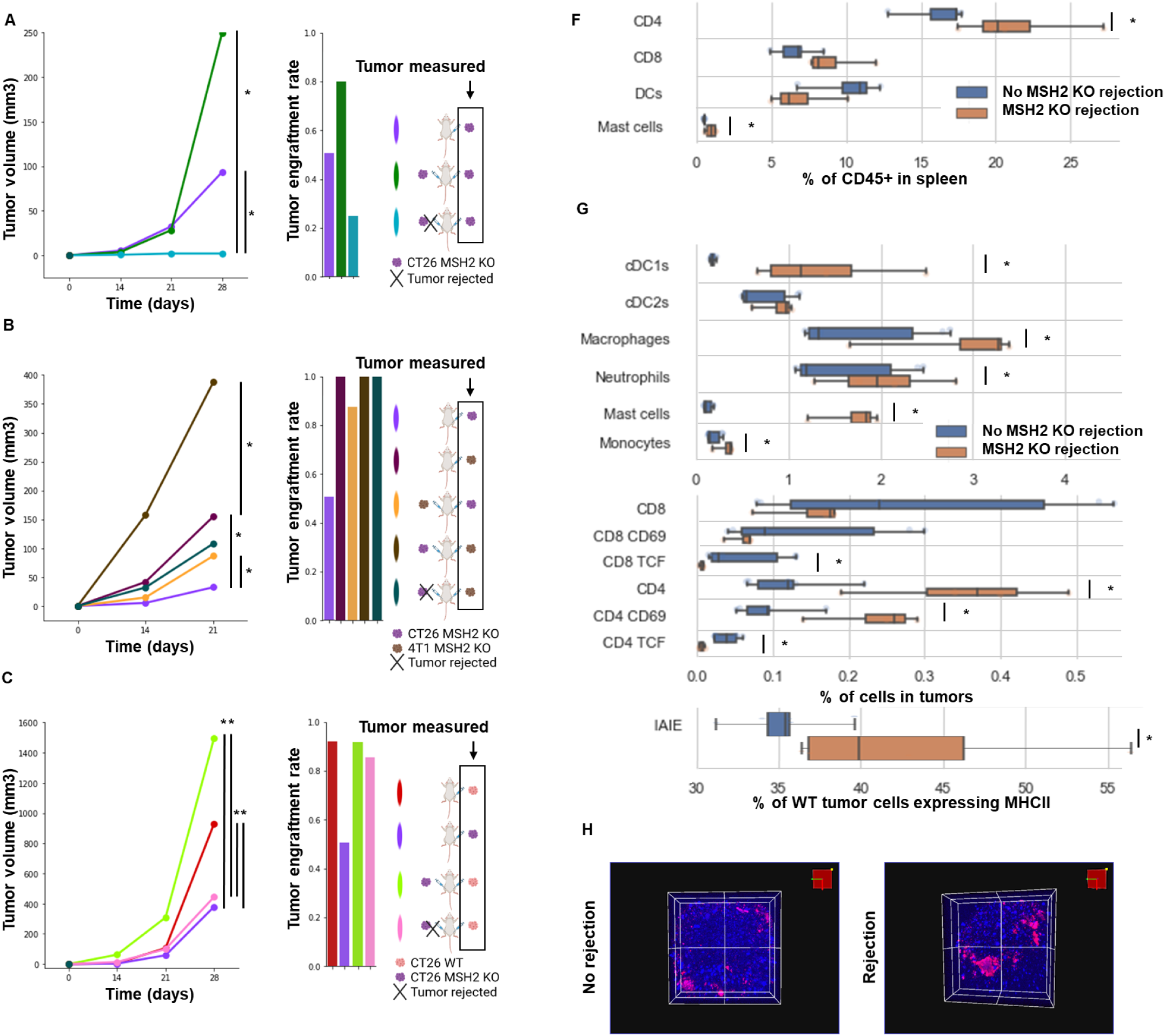
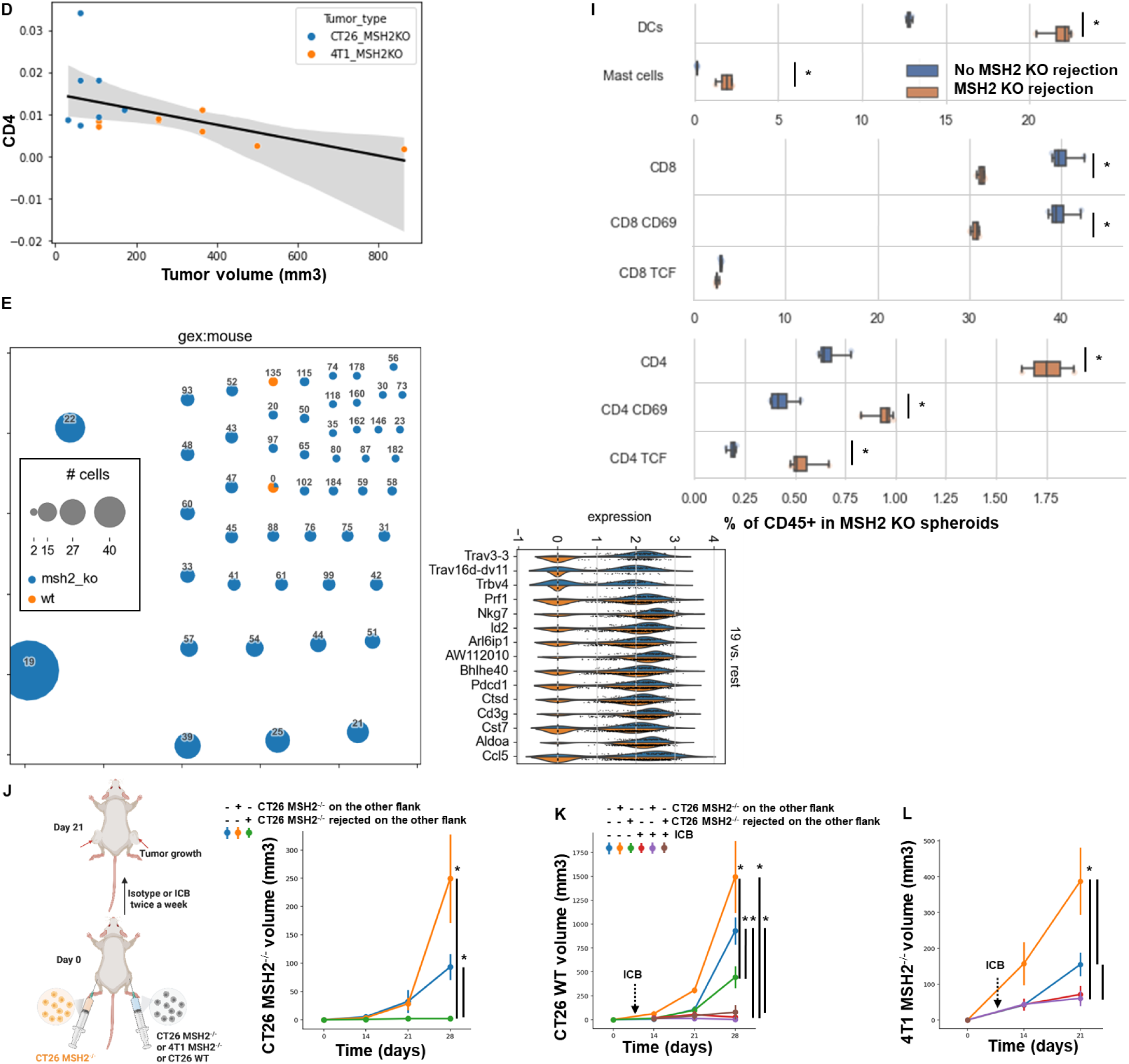
Tumor volume and immune infiltration of CT26 and 4T1 MMRd and MMRp tumors. **A/B/C**Tumor volume and engraftment in vivo over 28 days (200,000 cells/tumor). N= 6 to 27, Mann-Whitney U Test, pvalue<0.05. **A** Tumor volume of CT26 MSH2 KO tumors according to the presence of a CT26 MSH2 KO tumor on the other flank. **B** Tumor volume of CT26 MSH2 KO tumors according to the presence of a 4T1 MSH2 KO tumor on the other flank. Tumor volume of 4T1 MSH2 KO tumors according to the presence of a CT26 MSH2 KO tumor on the other flank. **C** Tumor volume of CT26 WT tumors according to the presence of a CT26 MSH2 KO tumor on the other flank. **D** Infiltration of CT26 and 4T1 MSH2KO tumors by immune cells on mice with a CT26 MSH2 KO tumor an the left flank and a 4T1 MSH2 KO tumor on the right flank (flow cytometry) N= 7 to 8, Linear regression, pvalue<0.05. **E** scRNAseq and TCRseq clustering on immune cells infiltrating tumors expressing TCRs in WT or MSH2 KO tumors. The dataset contains only α/β T-cell receptors. Filtering to exclude all cells that don’t have at least one full pair of receptor sequences and multichain cells. Each dot represents a clonotype, e.g. cells with identical receptor configurations. The size of each dot refers to the number of cells with the same receptor configurations. Overexpressed genes in the more abundant clonotype (clonotype 19). **F** Spleen composition from mice that rejected or not a CT26 MSH2 KO tumor. **G** Infiltration of CT26 WT tumors by immune cells from mice that rejected or not a MSH2 KO CT26 tumor. **H** Infiltration of MSH2 KO spheroids by immune cells from mice that rejected or not a MSH2 KO tumor. **I** Infiltration of MSH2 KO CT26 spheroids by immune cells from mice that rejected or not a MSH2 KO tumor. **J/K/L** Tumor volume in vivo over 28 days (200,000 cells/tumor). Unilateral or bilateral injections. N= 4 to 27, Mann-Whitney U Test, pvalue<0.05. Data are represented as mean ± SEM. **J** Tumor volume of CT26 MMRd tumors according to the presence of a CT26 MMRd tumor on the other flank. **K** Tumor volume of CT26 WT tumors according to the presence of a CT26 MMRd tumor on the other flank and ICB (anti-PD-1 + anti-CTLA4 + anti-LAG3 + anti-TREM2 twice a week after day 7, 100 ug each). Data are represented as mean ± SEM. **L** Tumor volume of 4T1 MMRd tumors according to the presence of a CT26 MMRd tumor on the other flank and ICB (anti-PD-1 + anti-CTLA4 + anti-LAG3 + anti-TREM2 twice a week after day 7, 100 ug each) (*8*). Data are represented as mean ± SEM.

**Extended Figure 4.**
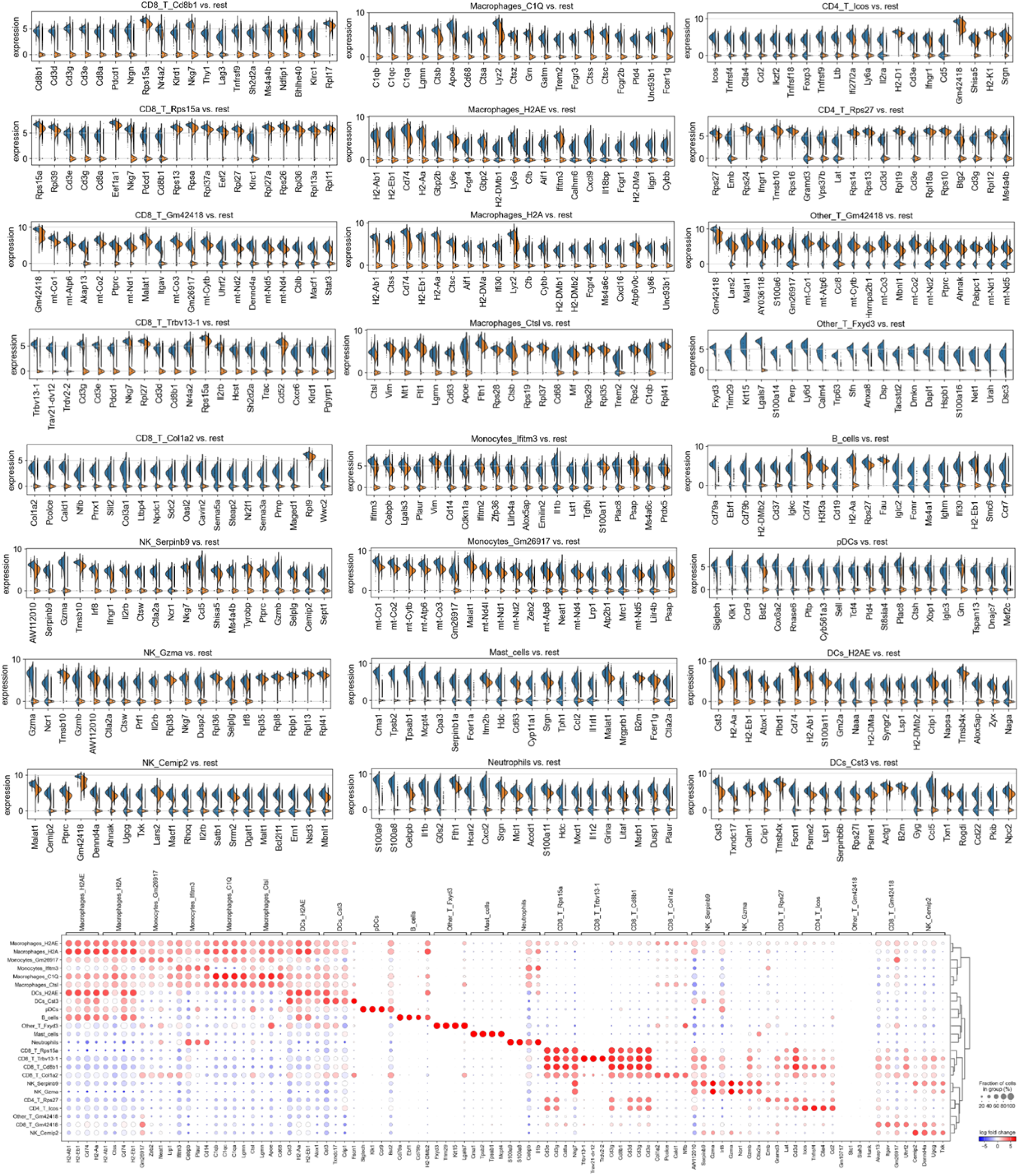
Single cell analysis of immune cells in CT26 MSH2 KO tumor with or without vaccination and anti-PD-1 therapy. ScRNASeq of immune cells in CT26 MSH2 KO tumor with or without vaccination and anti-PD-1 therapy. More significantly upregulated genes in each immune cluster (Wilcoxon test). N=3 tumors per condition.

**Extended Figure 5.**
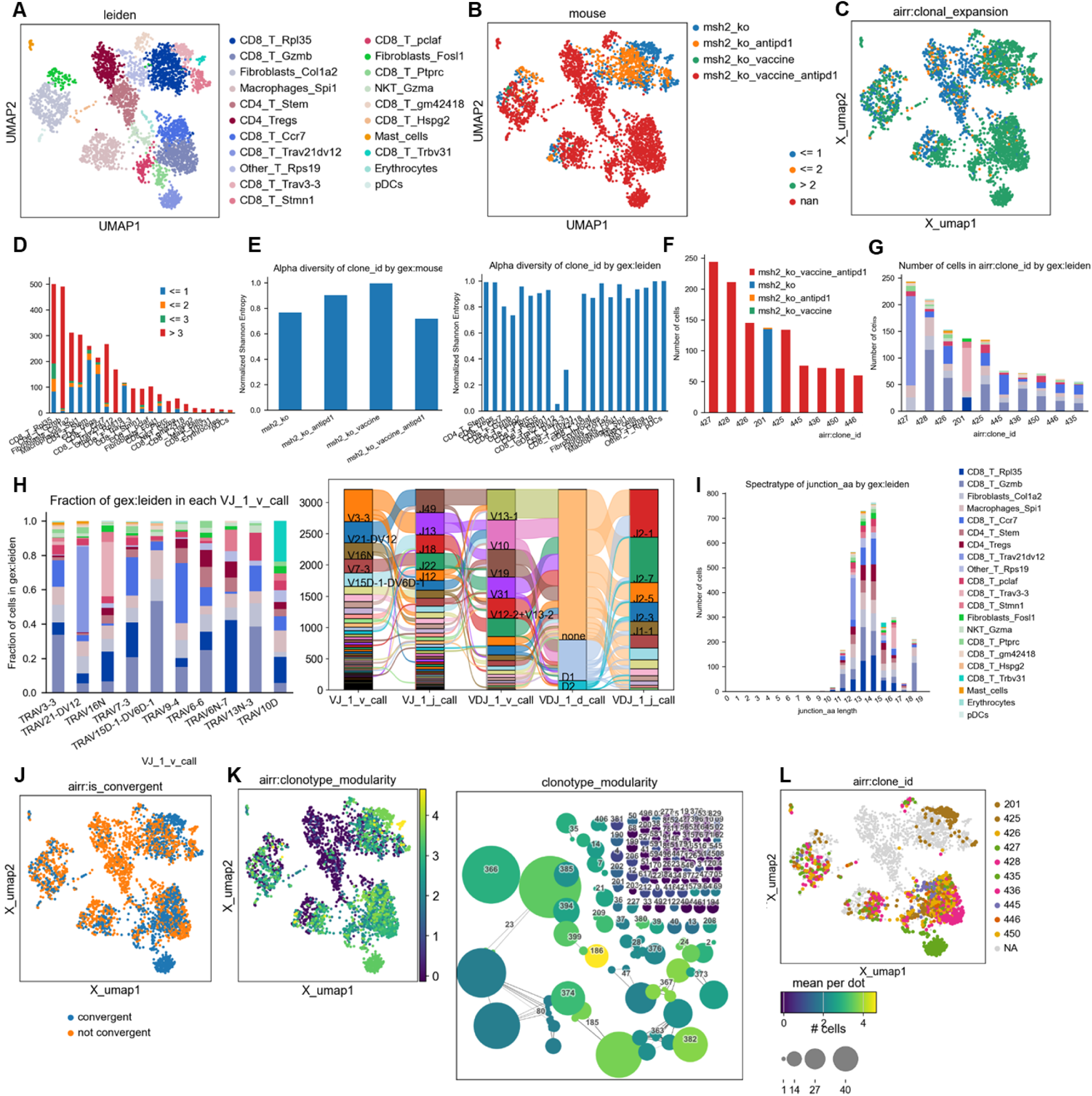
TCRseq analysis paired with scRNAseq analysis on CT26 MSH2 KO tumor according to vaccination and anti-PD-1 therapy. **A/B** scRNAseq clustering on immune cells infiltrating tumors expressing TCRs. The dataset contains only α/β T-cell receptors. Filtering to exclude all cells that don’t have at least one full pair of receptor sequences and multichain cells. **C/D** Clonal expansion, number of cells or percentage for each clonotype. **E** Clonal expansion and diversity. **F/G** Clonotype abundance in each cluster and in each condition. **H** Most abundant V-genes and exact combinations of VDJ genes. **I** Spectratype and length distribution of CDR3 regions. **J** Convergent evolution. **K** Clonotype modularity. **L** UMAP of the most abundant clones. Analysis was performed using scirpy and scanpy.

**Extended Figure 6.**
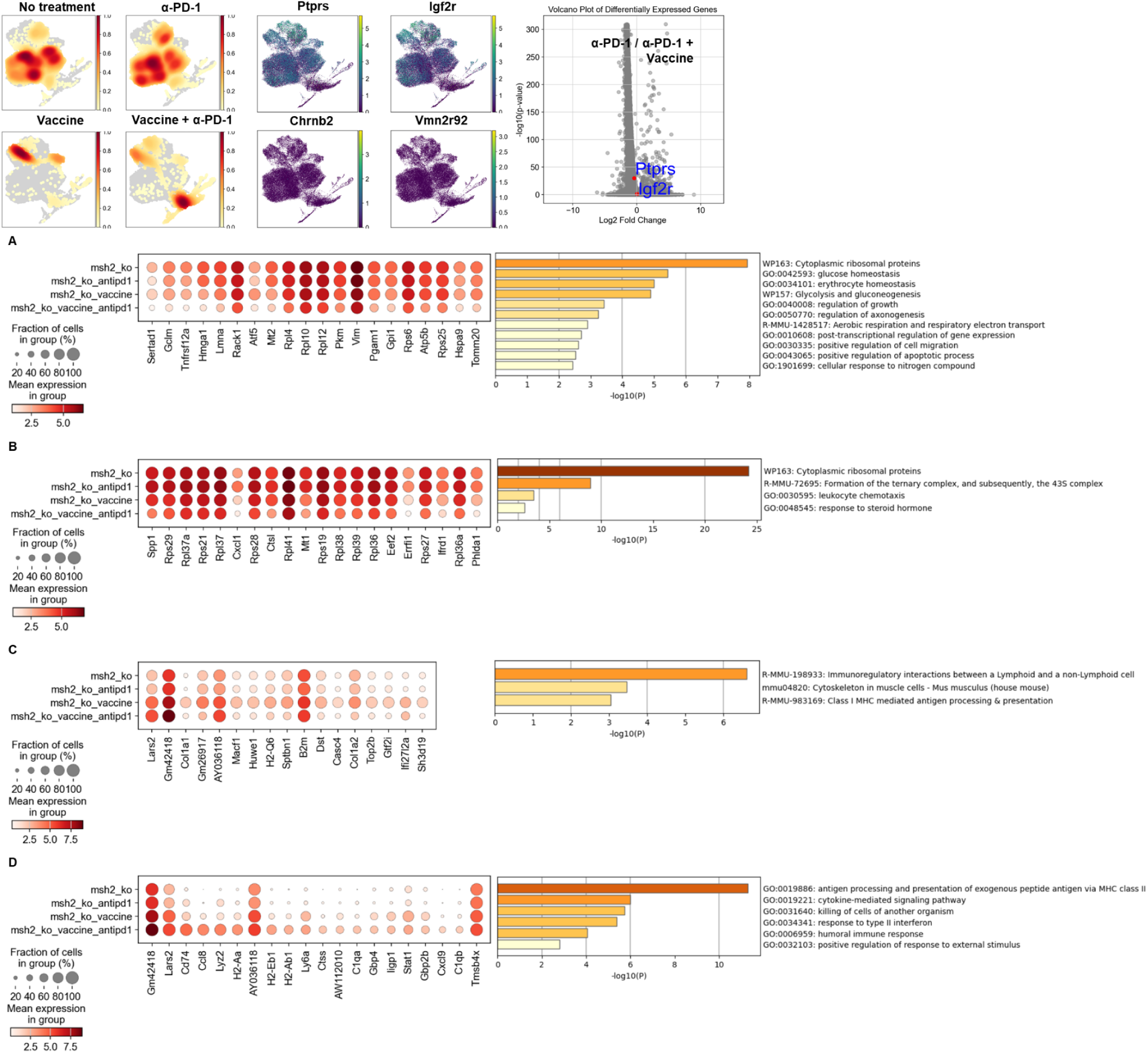
Single cell analysis of tumor cells in CT26 MSH2 KO tumor with or without vaccination and anti-PD-1 therapy. ScRNASeq of tumor cells in CT26 MSH2 KO tumor with or without vaccination and anti-PD-1 therapy. More significantly upregulated genes in each condition (Wilcoxon test), and gene enrichment analysis by metascape. N=3 tumors per condition. **A** Upregulated genes and pathways in untreated tumor cells. **B** Upregulated genes and pathways in tumor cells following PD-1 blockade. **C** Upregulated genes and pathways in tumor cells following vaccination. **D** Upregulated genes and pathways in tumor cells following PD-1 blockade + vaccination.

